# A resource for analyzing *C. elegans*’ gene expression data using transcriptional gene modules and module-weighted annotations

**DOI:** 10.1101/678482

**Authors:** Michael Cary, Katie Podshivalova, Cynthia Kenyon

## Abstract

Identification of gene co-expression patterns (gene modules) is widely used for grouping functionally-related genes during transcriptomic data analysis. An organism-wide atlas of high quality fundamental gene modules would provide a powerful tool for unbiased detection of biological signals from gene expression data. Here, using a method of independent component analysis we call DEXICA, we have defined and optimized 209 modules that broadly represent transcriptional wiring of the key experimental organism *C. elegans*. Interrogation of these modules reveals processes that are activated in long-lived mutants in cases where traditional analyses of differentially-expressed genes fail to do so. Using this resource, users can easily identify active modules in their gene expression data and access detailed descriptions of each module. Additionally, we show that modules can inform the strength of the association between a gene and an annotation (e.g. GO term). Analysis of “module-weighted annotations” improves on several aspects of traditional annotation-enrichment tests and can aid in functional interpretation of poorly annotated genes. Interactive access to the resource is provided at http://genemodules.org/.

## Introduction

Nearly half of the predicted protein-coding genes in *Caenorhabditis elegans* lack a functional annotation based on direct experimental evidence^1^. As a result, querying gene annotation databases, such as the Gene Ontology (GO) or curated-pathways can fail to detect biologically meaningful signals in gene expression data ^2–4^. An alternative approach to understanding gene function is to use information about genes’ transcriptional activity. Gene-expression data can be used to define groups of genes that show similar patterns of expression, or co-variation, across multiple conditions. These groups are called transcriptional gene modules, with each module potentially representing a discrete biological phenomenon. Gene modules are routinely constructed when clustering algorithms are applied to gene expression data and have been used successfully to identify gene regulatory mechanisms in a variety of contexts, from the yeast cell cycle^5^ and sporulation^6^ to human cancer cells^7^ to cognitive decline in patients with Alzheimer disease^8^. Furthermore, large compendia of data sampling diverse perturbations have been used to define fundamental gene-expression programs of entire organisms^9–11^.

In *C. elegans*, the most recent effort to generate high-quality fundamental transcriptional modules is now almost two decades old^12^. Co-expressed genes were grouped together into 43 groups (or “mountains”) based on their correlation across 553 microarrays. However, the compendium did not contain all genes (e.g. over 30% of the microarrays only contained ∼11,000 of *C. elegans’* 20,470 protein-coding genes) and each gene was assigned exclusively to one group, although it is well-established that genes can participate in multiple processes ^13, 14^. The number of perturbations for which gene expression data are available has also increased substantially since 2001.

Here, we define 209 transcriptional gene modules in *C. elegans* using a heterogeneous compendium of 1386 microarrays and a method we call DEXICA, for Deep EXtraction Independent Component Analysis. DEXICA builds on prior implementations ^10, 15, 16^ of independent component analysis (ICA) for gene module extraction by maximizing the biological information content of the modules. It does so by varying data pre-processing methods and the number of extracted ICA components (modules) until the number of biological annotations is maximized. DEXICA also uses an artificial neural network to partition each independent component, i.e. gene module, to determine which genes should be included and which should be excluded.

We show that the 209 DEXICA *C. elegans* modules capture gene expression patterns that correspond to biological processes; for example, responses to osmotic stress, xenobiotics and pathogenic bacteria, and to several individual tissues. Furthermore, data analysis in the module space correctly reveals biological processes that are missed by analyses of differentially-expressed genes. We provide a user-friendly web interface in which users can test which of the 209 gene modules are active in their datasets and find detailed information about each module that helps determine which biological process(es) they represent.

Finally, we explore whether gene modules can be used to improve existing gene annotations. We reason that an annotation shared among co-expressed genes is more likely to be relevant to their function than one that is not, and that annotations of co-expressed genes can provisionally be “transferred” onto their poorly-annotated companions. We calculate what we call “module-weighted gene annotations” by weighting the association between a gene and an annotation by the degree to which the annotation appears predictive of the gene’s module membership. We show that matrix-based analysis of module-weighted annotations is more sensitive and specific than common annotation enrichment tests. We provide a framework for using module-weighted annotations to detect significant GO terms and promoter oligonucleotides directly from expression data, and to identify novel GO terms conferred onto genes based on their module membership.

## Results

### Development of DEXICA and extraction of gene modules

A large body of gene expression data is publicly available^17, 18^ and has enabled computational prediction of gene modules^10, 12, 19–21^. We refer to our method for going from a raw compendium of gene expression data to an optimized set of gene modules and a list of genes that belong to each module as DEXICA, for <u>D</u>eep <u>EX</u>traction Independent <u>C</u>omponent <u>A</u>nalysis (described below).

While several methods exist for defining gene modules, independent component analysis (ICA) generally outperforms clustering-based approaches and principle component analysis in extracting biologically relevant signals from large datasets^15, 22–24^. Advantages of ICA include its ability to deal well with high dimensional data and to generate modules that can share genes. Furthermore, ICA does not assume that latent signals in the data follow a Gaussian distribution, an important property for gene module prediction, as gene regulation signals appear to be primarily super-Gaussian^25^. For these reasons, we chose to use ICA for module construction.

Briefly, ICA is a blind source separation method that attempts to “unmix” a signal comprising additive subcomponents by separating it into statistically-independent source signals^26^. In the common notation, an *n* × *m* data matrix, ***X***, is decomposed into two new matrices – a *n* × *k* source matrix, ***S***, and an *k* × *m* mixing matrix, ***A***, where *k* is the number of independent components:

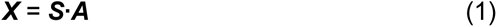

In the context of gene-expression analysis, ***X*** is a matrix of *m* measurements (e.g. microarrays) of *n* genes, and *k* independent components are interpreted as gene modules. ***A*** indicates the weight of each module in each microarray and ***S*** indicates the relative level of inclusion of each gene in each module^27^ (Figure 1a).

**Figure 1.**
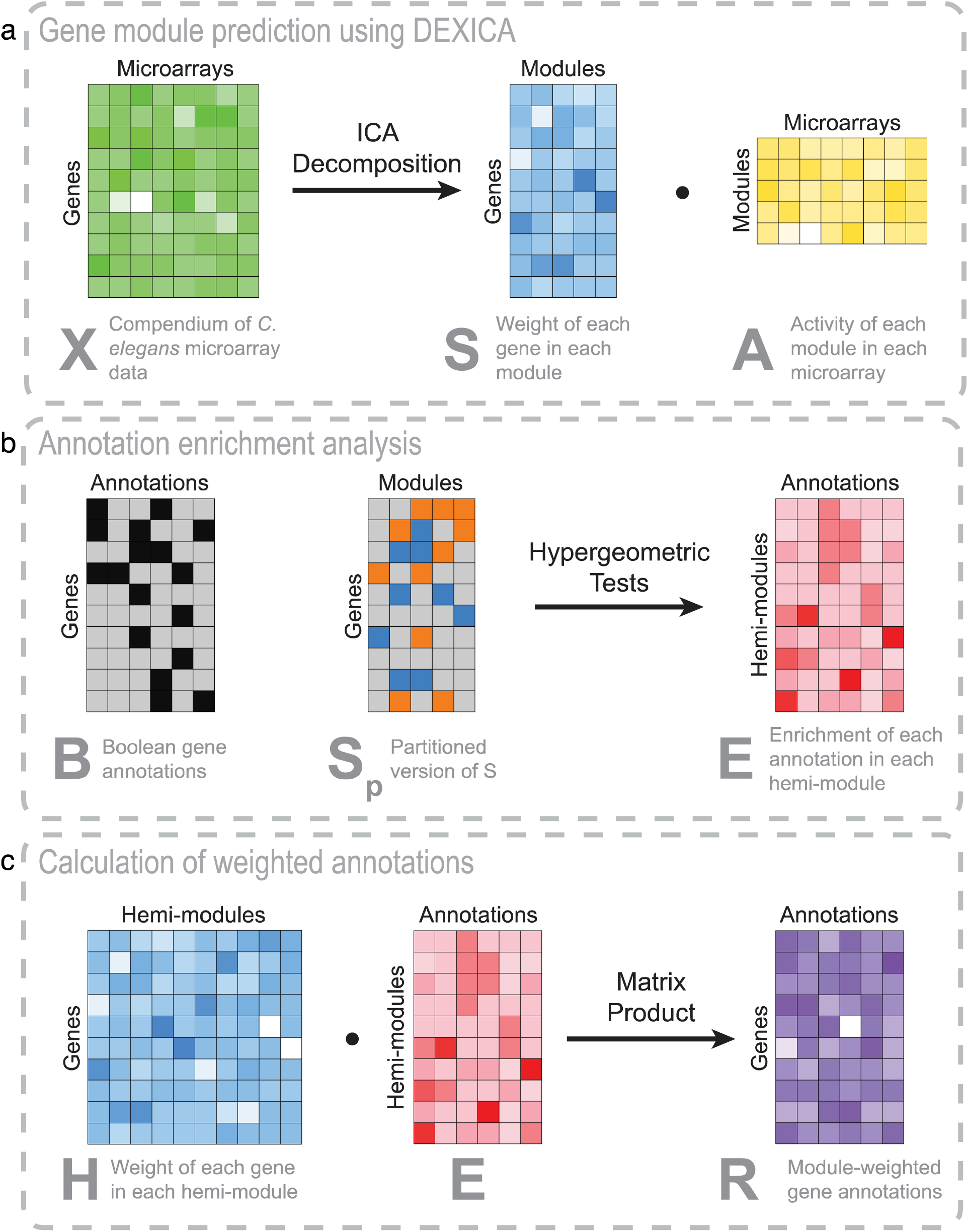
Schematic of module prediction using DEXICA and derivation of module-weighted annotations. (**A**) A matrix of gene expression data, ***X***, is decomposed using independent component analysis (ICA) into a gene module definition matrix, ***S***, and a matrix containing the weight of each module in each microarray, ***A***. Rows of the ***S*** matrix indicate the relative degree of inclusion of a given gene in each module. (**B**) Annotation enrichment within modules is calculated using matrices ***B*** and ***S_p_***. Known associations between genes and annotations are captured in a Boolean matrix ***B***, where 1 indicates an association between a gene and an annotation and 0 indicates a lack of an association. ***S*** is partitioned into ***S_p_***, where 0 indicates a gene’s exclusion from a module, while 1 indicates a gene’s assignment to the positive hemi-module and −1 to the negative hemi-module (a hemi-module comprises genes that have extreme weights and the same sign in a given column of the ***S*** matrix). Enrichment of genes associated with each annotation in each hemi-module is calculated using hypergeometric tests and log(p-values) are recorded in a new matrix, ***E***. (**C**) To generate module-weighted annotations, the ***S*** matrix is first transformed into a matrix, ***H***, which indicates the weight of each gene in each hemi-module. ***H*** contains twice as many columns as ***S*** because there are two hemi-modules per module. The dot product between ***H*** and ***E*** results in the ***R*** matrix, where a row indicates the weighted association between a gene and each annotation. The color intensity in ***R*** indicates annotation weights that would result given the indicated expression values in ***X*** and the Boolean annotations in ***B***. Color saturation in all matrices except ***B*** (Boolean) and ***S_p_*** (trivalued) indicate relative numeric values, with least saturated colors (i.e., white) indicating highly negative values, and most saturated colors indicating highly positive values.

Using simulated data, we found that module-prediction accuracy was highest when the number of extracted components matched the true number of modules (Supplementary Figure 1). Therefore, we sought to optimize module prediction by evaluating results based upon their biological information content, such as enrichment of Gene Ontology (GO) terms^2^, REACTOME pathways^4^, and tissue-specific expression^28^. We applied ICA to a diverse compendium of 1386 *C. elegans* Affymetrix arrays obtained from the Gene Expression Omnibus (GEO) database^18^. We then compared results obtained from a wide variety of preprocessing methodologies and total number of components extracted. Omitting between-experiment quantile normalization from the preprocessing procedure produced modules that were more annotation rich than those produced by a published implementation of ICA-based module extraction ^10^ (Figure 2a-c). Furthermore, annotation content of the modules was maximized when the number of gene modules (i.e. independent components) ranged from 191 to 226.

**Figure 2.**
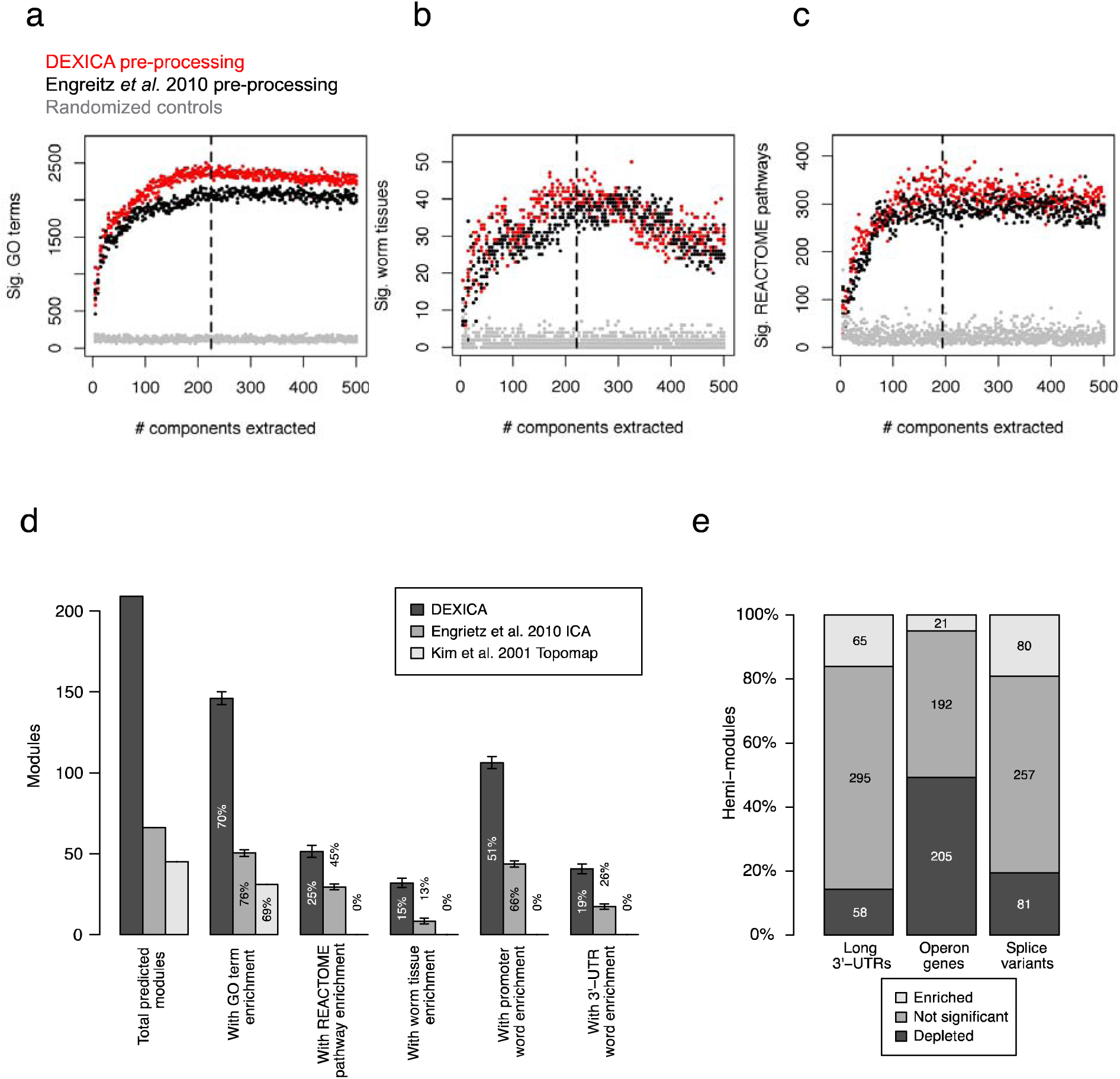
Optimization of gene modules. To determine the optimal preprocessing method and the optimal number of components (gene modules) to extract from a gene expression compendium of *C. elegans* microarray data, we calculated the number of Gene Ontology terms (**A**), *C. elegans* tissues (**B**), and REACTOME pathways (**C**) that were significantly enriched in at least one gene module. Black points show results from a compendium produced using a preprocessing procedure used by Engreitz *et al.*^10^; red points show results for the best alternative preprocessing method that we tested. Black dashed lines indicate the point on the x-axis of each graph at which loess regression curves showed the greatest difference between red points and results from randomized controls (grey points). (**D**) The number modules produced by DEXICA, by a different ICA-based method used by Engreitz *et al.*^10^, and by *C. elegans* gene expression topomap generated by Kim *et al*.^12^. The number and the percentage of total predicted modules that have significant enrichment for various annotations are shown. Error bars indicate s.d. between repeat runs of DEXICA or Engreitz *et al*. method. (**E**) Each of the 418 hemi-modules was tested for enrichment of genes with different structural properties – presence a long 3’-UTR, transcription as part of an operon, and presence of multiple annotated splice variants.

To enable functional evaluation of the modules, it is useful to summarize them as discrete gene sets. To this end, we partitioned each column of the ***S*** matrix into three sets of genes: one set consisting of genes excluded from the module, and two other sets consisting of genes regulated in opposite directions. We refer to these latter two sets as “hemi-modules”, one set consisting of genes with highly positive weights and another consisting of genes with highly negative weights (signs assigned arbitrarily) in the independent component. While others have used a fixed-threshold approach to component partitioning^10, 29, 30^; for example, defining genes with weights exceeding +/-3 standard deviations from the component mean to be “in” each hemi-module, we found that individual modules showed maximum annotation enrichment at different thresholds, suggesting that a ‘one-size-fits-all’ approach to partitioning is sub-optimal. An alternative approach to partitioning that we attempted (described in Frigyesi *et al.*^31^) failed to converge in many cases (data not shown). Therefore, to increase partitioning accuracy, we trained a function to predict partitioning thresholds from the shape of component distributions. Because plots of training data revealed a complex solution surface, we decided to use an artificial neural network (ANN) to predict partitioning thresholds for each component from the skewness and kurtosis of its distribution. The output of the ANN, i.e. predicted partitioning thresholds, is shown in Supplementary Figure 2a. Although using a fixed-threshold approach to module partitioning produced similar results qualitatively (Supplementary Figures 2b-d), it resulted in fewer significant annotations across the range of parameters tested than did ANN-based partitioning (p < 2.2E-16, Supplementary Figure 2e). Because the mean optimum number of extracted components (dashed vertical lines in Figure 2a-c and Supplementary Figures 2b-d) was similar for both threshold and ANN partitioning (209, and 209.33, respectively), we chose 209 as the final number of components to extract from the compendium.

Gene modules are expected to represent sets of genes that are co-regulated at the level of mRNA expression or stability. Therefore, DEXICA modules should be enriched for DNA regulatory sequences. To test this, we generated a list of potential regulatory oligonucleotide sequences (called ‘words’) by applying the Mobydick algorithm^32^ to the set of all predicted *C. elegans* promoter regions and, separately, to the set of all predicted *C. elegans* 3’-UTRs. We then calculated the statistical significance of the over- or under-representation of genes bearing each word in each gene module. Across multiple runs with 209 components, the mean number of gene modules containing significant promoter words and 3’-UTR words was 106.3 and 40.6, respectively, significantly greater than results produced by other module prediction methods we tested (p < 2.2E-16, Figure 2d).

Because the ICA algorithm employed by DEXICA, *fastICA*, converges to a final solution from a random starting point^26^, small differences typically exist in the output of different runs; these differences can be seen in the vertical spread of data points in Figures 2a-c, and in the error bars in Figure 2d. While others have reconciled such differences through a clustering approach applied to the output of numerous runs of the algorithm (so called “iterated ICA”)^10, 31^, when applied to the *C. elegans* Affymetrix compendium, many of the final components generated by this method were highly correlated to one another, indicating non-independence and potential redundancy among the components (data not shown). We therefore sought to choose a single, high quality, *fastICA* run output to use as predicted gene modules. Because we considered word enrichment the most unbiased measure of module quality (as it relies only on DNA sequence data), we chose as our final module set (Supplementary Table 1) the run from a set of 100 with the best combined rank of significant promoter and 3’-UTR words (it ranked first in promoter words and third in 3’-UTR words, Supplementary Figure 3). The mean module size is 385 genes, with the majority of modules having fewer than 385 genes (the smallest module contains 49 genes and the largest contains 2383 genes).

Co-expressed genes often share common structural elements. For example, genes within operons are switched on together during recovery from growth-arrested states in *C. elegans*^33^ and 3’-UTR length is associated with proliferation in cancer cells^34^. To further explore the information content of DEXICA-extracted modules, we tested each hemi-module for over- and under-enrichment of genes with long 3’UTRs, for genes appearing in operons and for genes with multiple splice forms. Of the 418 hemi-modules, 65 contained a significant bias toward long 3’-UTR genes and 58 contained a bias toward short 3’-UTR genes (q < 0.1, threshold chosen based on randomized control trials, see below; Figure 2e). Twenty-one hemi-modules were significantly enriched and 205 hemi-modules were significantly depleted for operon genes, and 81 hemi-modules were enriched and 80 hemi-modules were depleted for genes with multiple splice variants (Figure 2e). Control tests performed on the same module set but with randomly scrambled gene IDs produced no significant modules below q = 0.1 for any of the gene properties we tested (data not shown). Therefore, DEXICA successfully groups genes with common structural features into modules, consistent with the known relationship between gene structure and expression.

Together, these results show that the 209 *C. elegans* gene modules extracted from a large microarray compendium are enriched for GO terms, tissue and pathway annotations as well as for potential regulatory DNA sequences and gene structural properties. DEXICA is available as an R package (https://github.com/MPCary/DEXICA). It provides tools to optimize ICA module extraction and partitioning based on annotation enrichment and can be applied to any gene expression compendium.

### *C. elegans* DEXICA modules are biologically informative

We observed enrichment of functional annotations and oligonucleotide sequences in DEXICA-extracted modules with even the least optimal parameter settings (see Figure 2). To further test the biological significance of the final 209 modules and to begin annotating them, we constructed an alternate microarray compendium comprising not 1386 individual arrays, as in the original compendium, but rather the gene fold changes arising from contrasting experimental and control samples in the same experiment. Projecting the resulting 716-column matrix (one for each contrast) into the space defined by our gene modules allowed us to see which experimental perturbations activate or inhibit which modules.

We compared the most enriched GO terms in each module to the most strongly activating or inhibiting experimental perturbations and observed that in many cases these were in obvious agreement (Supplementary Table 2). For example, the strongest activator of m10 (module 10) in the compendium is a mutation in *osm-7*, a gene needed to respond to hypertonic stress through accumulation of glycerol. Consistent with this, the top GO terms enriched within genes that comprise m10 describe glycerol metabolism and accumulation. Module 153 is strongly activated by mutations in *lin-35*, a gene that is part of the DRM/DREAM complex, and accordingly, this module is also enriched for “DRM complex” GO terms. Similarly, m200 is activated in response to the pathogen *P. aeruginosa* and the top GO terms enriched within m200 genes include “defense response to Gram-negative bacteria” and “innate immune response”.

DEXICA modules can also capture gene expression patterns that distinguish different tissues. For example, m23 is strongly activated by neuronal RNA and the top GO term enriched in this module is “neuropeptide signaling”. Similarly, m144 distinguishes muscle cells from other cell types in the worm and is enriched for the “sarcomere organization” and “striated muscle dense body” GO terms. Together, these results indicate that DEXICA modules represent biologically meaningful sets of genes.

It is important to note that a lack of an obvious agreement between the nature of a perturbation that strongly activates a module and the top GO terms enriched in that module does not necessarily indicate that a module is biologically meaningless. On the contrary, this apparent disagreement can be a powerful tool for understanding complex transcriptional responses to perturbations. For example, starvation of L1 larvae induces a developmental arrest and has widespread effects on metabolism^35^. Module 51 is strongly activated by starvation and is enriched for “snoRNA/nucleolus” GO terms.

Thus, this module represents genes that play a role in ribosome biogenesis and protein synthesis, reduction of which is an important response to starvation. Obvious agreement between a perturbation and GO terms would also be absent if a process had not previously been associated with a particular condition, making modules useful for hypothesis generation. For example, a wild *C. elegans* isolate, JU1580, shows strong activity of m55 relative to the laboratory *C. elegans* strain, N2, (Supplementary Table 2) and genes within m55 are enriched for the “innate immune response” GO term. This suggests that the immune systems of wild and lab *C. elegans* strains have different levels of basal activity, a connection that, to our knowledge, has not been proposed previously.

### Annotation of DEXICA modules

Detailed information about modules that are active in a gene-expression experiment would be of primary interest to an investigator. As a comprehensive resource, we created a Module Annotation Page for each of the 209 modules (see Data availability, Tool 3). Each page shows the module’s significantly enriched and depleted GO terms (excerpted in Supplementary Table 2), REACTOME pathways, tissues of known expression, enriched promoter and 3-’UTR oligonucleotide words, and component genes (each linked to a WormBase description).

A limitation of using GO terms and other pre-existing annotations to interpret gene modules is that some cellular activities are poorly annotated (see UPR^mt^ example below). For this reason, an additional strategy to understanding what a given module represents is to examine its activity under a variety of conditions. To facilitate this, in the Module Annotation Pages we have also provided a ranked list of perturbations (from the 716 we generated) that activate each module significantly (also see Supplementary Table 2). While it is unlikely that a typical gene module can be described completely using a single semantic annotation due to the complexity of gene regulatory networks, we suggest processes that modules likely represent based on our comparison of module activity under several conditions and our examination of enriched annotations (Supplementary Table 2). In a typical workflow, once the active modules in a gene expression experiment are identified by a researcher, we recommend that the researcher consult Module Annotation Pages and Supplementary Table 2 to infer the biological process(es) described by the modules of interest.

### DEXICA modules reveal processes missed by conventional tools

#### Mitochondrial unfolded protein response

To test the utility of DEXICA modules for interpreting *C. elegans* gene expression data, we asked whether we could identify known biological processes activated in a mutant with a complex phenotype. Mutations in components of the electron transport chain reduce respiration, slow development and reproduction and increase lifespan in *C. elegans*^36^. The mitochondrial unfolded protein response (UPR^mt^) is induced in response to a stoichiometric imbalance between nuclear and mitochondrial proteins within mitochondria, and this process is known to be active in long-lived respiration mutants^37, 38^. However, analysis of significantly differentially expressed genes in the *isp-1* respiration mutant, either using GO terms or KEGG pathways, failed to identify UPR^mt^ or, for that matter, any process related to mitochondria or a response to stress (Figure 3a).

**Figure 3.**
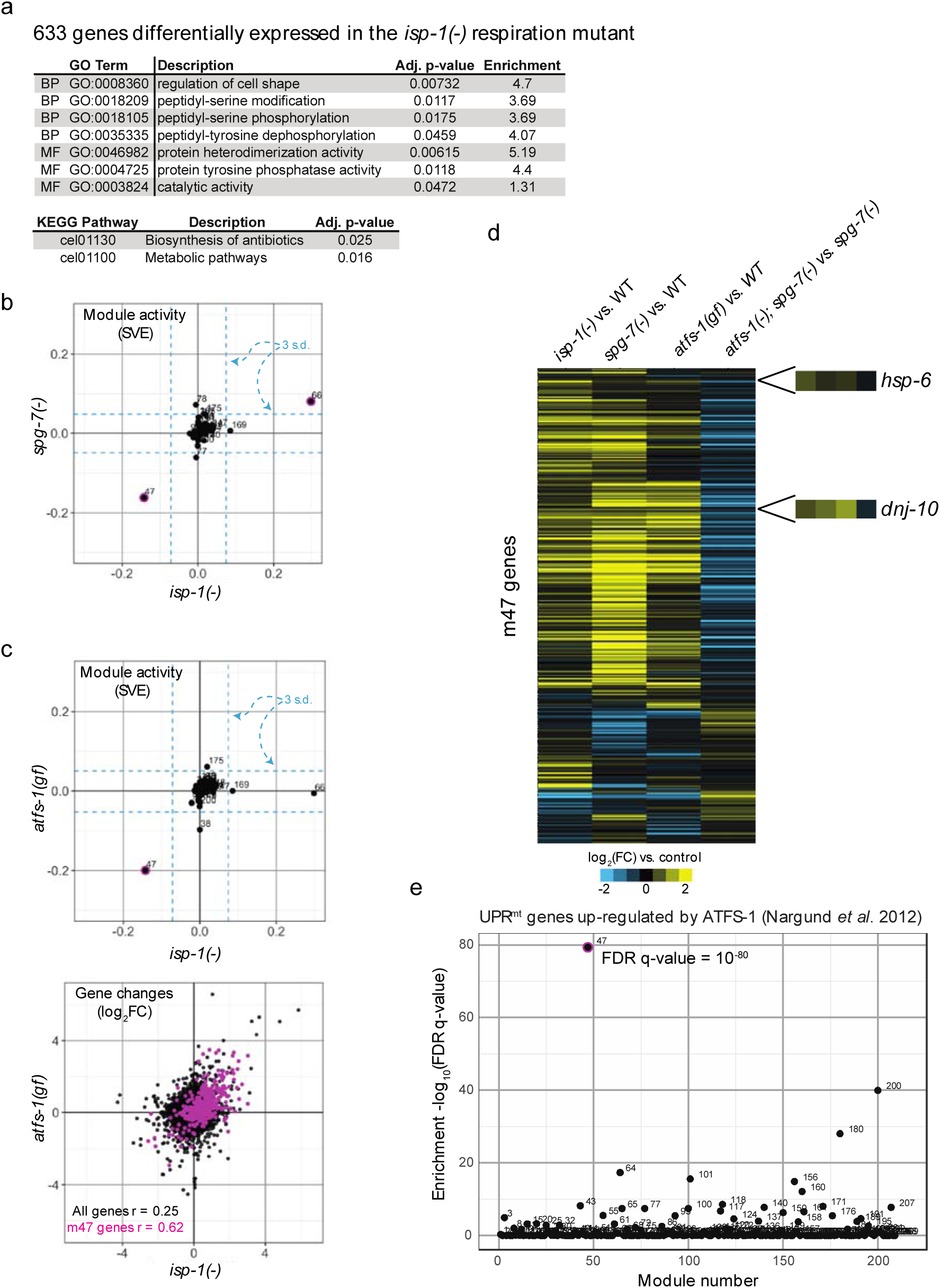
Analysis of module activity in *isp-1* respiration mutants reveals activity of the key mitochondrial unfolded protein response (UPR^mt^) transcription factor, ATFS-1. (**A**) Enrichment of GO terms and KEGG pathways within genes differentially expressed in *isp-1(qm150)* respiration mutants. Significant gene expression changes were determined using the *limma* R package and FDR-adjusted p-value of < 0.05. (**B**) Knockdown of a protein quality-control protease *spg-7* is the top non-respiration microarray experiment in the compendium that induces activity of the *isp-1* module m47. Module activity is expressed as signed variance explained (SVE). (**C**) Comparison of *isp-1(qm150)* mutants and animals that express a constitutively-active form of *atfs-1*, a key transcription factor required for the UPR^mt^. The gene expression scatter plot shows all detected genes (22625; black) overlaid with genes that belong to m47 (367; pink). r, Pearson correlation coefficients. (**D**) Normalized changes in expression of genes that belong to m47 (367 genes) in response to inhibition of respiration (*isp-1* mutation, column 1), activation of UPR^mt^ by disrupting mitochondrial protein quality control (column 2), constitutive activation of *atfs-1* under normal conditions (column 3), and activation of UPR^mt^ in the absence of *atfs-1* (column 4). (**E**) Genes up-regulated by ATFS-1 during UPR^mt^, as determined in Nargund *et. al* 2012, are strongly enriched in m47. Enrichment was calculated using hypergeometric statistics.

Calculation of module activity (Tool 1) revealed that m47, m66 and m169 display the highest activity in the *isp-1* mutant (Figure 3b). These modules are strongly active in other respiratory chain mutants as well (see Module Annotation Pages), but the most strongly activating non-respiration perturbation of m47 is a mutation in *spg-7*, a mitochondrial protein quality-control protease, disruption of which is known to induce UPR^mt^ ^39^ (Figure 3b). This result suggested that m47 may encompass UPR^mt^ genes. If so, then m47 should be active in animals with constitutive activity of the transcriptional UPR^mt^ regulator, ATFS-1. To test this prediction, we calculated module activity in *atfs-1* gain-of-function mutants (this perturbation was not part of the original compendium, GEO Accession number GSE73669). Indeed, induction of the UPR^mt^ transcriptional response in otherwise healthy animals using this mutation strongly and specifically induced activity of m47 (Figure 3c). Furthermore, genes that belong to m47 showed concordant expression in the *isp-1* respiration mutant, in response to UPR^mt^ induction by disruption of *spg-7*, and in response to constitutive activity of ATFS-1 in normal animals (Figure 3c, d). As expected, mutation of *atfs-1* prevented these changes in animals with an induced UPR^mt^ (*spg-7* RNAi) (column 4 Figure 3d). Moreover, two chaperone genes known to be induced during UPR^mt^ (*hsp-6* and *dnj-10*) are part of m47. Taken together, these data strongly suggest that m47 represents genes induced by ATFS-1 during UPR^mt^. Therefore, analysis of module activity, but not GO term or KEGG pathway enrichment analyses, was able to correctly identify a key biological process using gene expression data from *isp-1* mutants.

Why did GO term enrichment analysis fail to recover the “mitochondrial unfolded protein response” term (GO:0034514)? To our surprise, there are only eight *C. elegans* genes annotated with this GO term (*haf-1*, *ubl-5*, *gcn-2*, *atfs-1*, *clpp-1*, *dve-1*, *hsp-6* and *hsp-60*). Among these, only *hsp-6* and *hsp-60* are induced during UPR^mt^, whereas the others are genes needed for activation of UPR^mt^. In the KEGG pathway database, UPR^mt^ is not included as a discrete entry, but is part of the “Longevity regulating pathway – worm” (map04212) entry, and similar to GO, primarily includes inducers rather than mediators of the UPR^mt^. These examples illustrate how incomplete annotations and/or overly general groupings (e.g., containing both activators and mediators) can result in a failure of standard annotation enrichment methods to detect biological signals in gene expression data.

Nargund *et al.* have defined a set of ATFS-1-targeted genes based on up-regulation of these genes with or without *atfs-1* and in the absence or presence of mitochondrial stress^40^. While module analysis was able to reveal UPR^mt^ without reliance on any pre-existing gene annotations or pre-defined gene sets, we wondered whether GSEA^41^ analysis of *isp-1* gene expression data using this gene set would have been able to identify ATFS-1 activity. The Nargund *et al.* ATFS-1 gene set showed enrichment within *isp-1* genes, but this enrichment was not statistically significant (Supplementary Figure 4). In contrast, when we tested enrichment of the ATFS-1 gene set within modules, we found a highly significant enrichment within m47 (Figure 3e). These results show that given a set of functionally related genes, testing enrichment of that set within modules can be more informative than testing enrichment within a ranked list of gene changes, likely because modules comprise groups of genes that are functionally related.

#### Hypoxia inducible factor 1

Reduction-of-function *isp-1* mutations extend lifespan in a manner dependent on the hypoxia inducible factor 1 (HIF-1)^42^, but, unexpectedly, we found that the overlap between the significant genes obtained by comparing *isp-1* or *hif-1* mutants to wild type was not statistically significant (*X^2^* test p-value = 0.17; Supplementary Figure 5a). The other highly-active modules in *isp-1* mutants besides m47 are m66 and m169. We wondered if these modules represent genes regulated by HIF-1. Indeed, when we compared gene module activity in *hif-1* and *isp-1* mutants, we found a strong anti-correlation (Pearson correlation between SVE = −0.730, p = 4.7E-36; Supplementary Figure 5b) driven by m66 and m169. The negative correlation is consistent with the finding that the life extension observed when *isp-1* activity is reduced requires activity of HIF-1. The idea that HIF-1 directly regulates transcription of genes in m66 and m169 is further supported by the fact that the canonical HIF-1 binding site [(A/G)CGTG] is the top oligonucleotide sequence enriched in the promoters of the genes comprising these modules (q-values 8.13e-105 and 1.47e-28, respectively). The similarity between module activity in *isp-1* and *hif-1* datasets, despite a lack of similarity among their most differentially expressed genes, suggests that the role of HIF-1 in regulating the lifespan of *isp-*1 mutants may be to instigate small but coordinated expression changes in many genes, most of which fail significance tests for differential expression in one or both datasets. These results demonstrate that gene modules are sensitive toward identification of transcriptional signatures and therefore can be useful for analysis of datasets where few gene changes reach statistical significance.

#### Gene modules as a hypothesis generation tool

As shown above, the correlation between the module activities of *hif-1* and *isp-1* was substantial (r = −0.730). To determine how often two sets of module activities generated from different experiments could be expected to show this degree of similarity, we determined module activity correlations for all possible pairs of the 716 experimental contrasts, excluding pairs in which both contrasts originated from the same experiment (i.e., from the same GEO series). This produced 188,805 contrast pairs, 13,376 (7.08%) of which showed a statistically significant correlation (Holm corrected p-value < 0.05). As expected, the highest-ranking pairs were the same experiment performed by different labs or at different times. The strength of the *hif-1* and *isp-1* correlation would have fallen within the top 1%, had those experiments been part of the contrast set. We include the full set of contrast comparisons in Supplementary Table 3, as gene expression changes that activate similar modules but that are generated by different experiments could prove useful to others for hypothesis generation. For example, had we observed the similarity between the *hif-1* and *isp-1* projections, we might have hypothesized a role for *hif-1* in *isp-1* mutants before such a role was discovered from genetic screening^42^.

### Using modules to improve gene annotations

Because gene annotations can be incomplete and biased, knowledge about which genes tend to be co-expressed in an organism can help inform which annotations are more likely to be biologically relevant and help infer annotations of orphan genes based on annotations of other module members (see Supplementary information for additional discussion). To this end, we devised a calculation of scores, which we call “module-weighted annotations”, based both on how much a given annotation is enriched in each module (matrix ***E***, Figure 1b-c) and the degree to which a given gene belongs to that same module (matrix ***H***, Figure 1c).

#### Module-weighted GO terms

To test the utility of module-weighted annotations, we first examined whether they generally recapitulate traditional (Boolean) annotations. If so, then genes traditionally associated with each annotation should have larger weights for those annotations (when normalization is omitted from their calculation) than do other genes (two-sample KS test, alpha level = 0.05). For this analysis we used GO terms as annotations and restricted the set to terms with at least 15 annotated genes in order to ensure robust signals; this set comprised 1651 GO terms. In 98.6% (1628/1651) of cases, genes associated with each term had significantly larger module-weighted annotations than did other genes, indicating that module-weighted GO annotations do recapitulate Boolean annotations.

We ranked each GO term by the weight of its most strongly associated gene (Table 1). GO terms with the most highly weighted genes were “ribosome” (CC) and “structural constituent of ribosome” (MF). Consistent with this finding, genes involved in ribosome biogenesis are known to be transcriptionally co-regulated^43^. On the other hand, GO terms associated with signal transduction and kinase activity showed the lowest gene weights. These results are statistically significant (see Online methods) and show that some types of gene groupings (e.g., genes encoding kinases) often used in the analysis of gene expression data may not actually represent functions that are coordinately regulated at the gene transcript level. Therefore, we suggest that researchers use caution when interpreting a result that certain annotations, like “kinases”, appear to be enriched in a transcriptomics experiment.

**Table 1.**
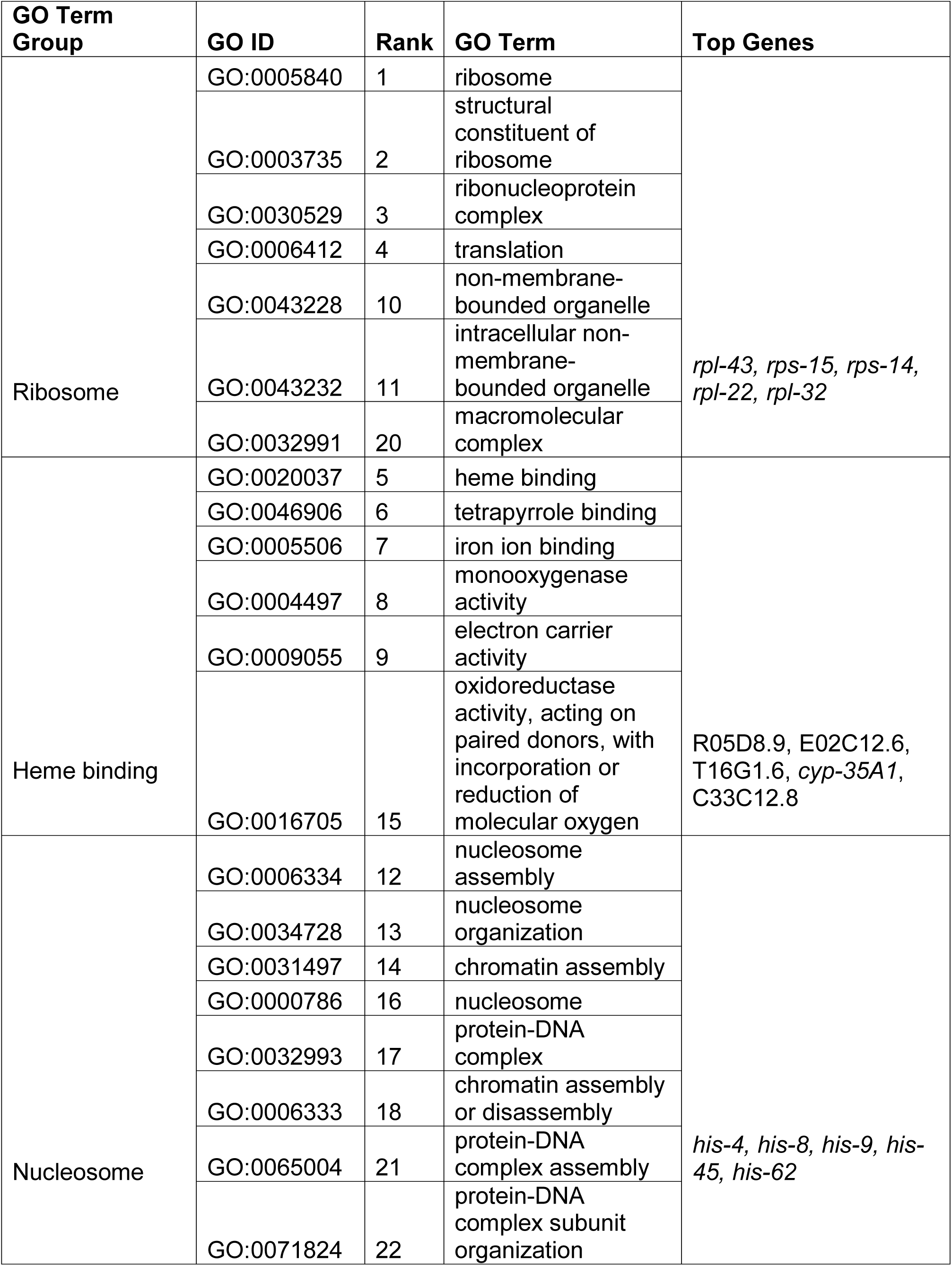
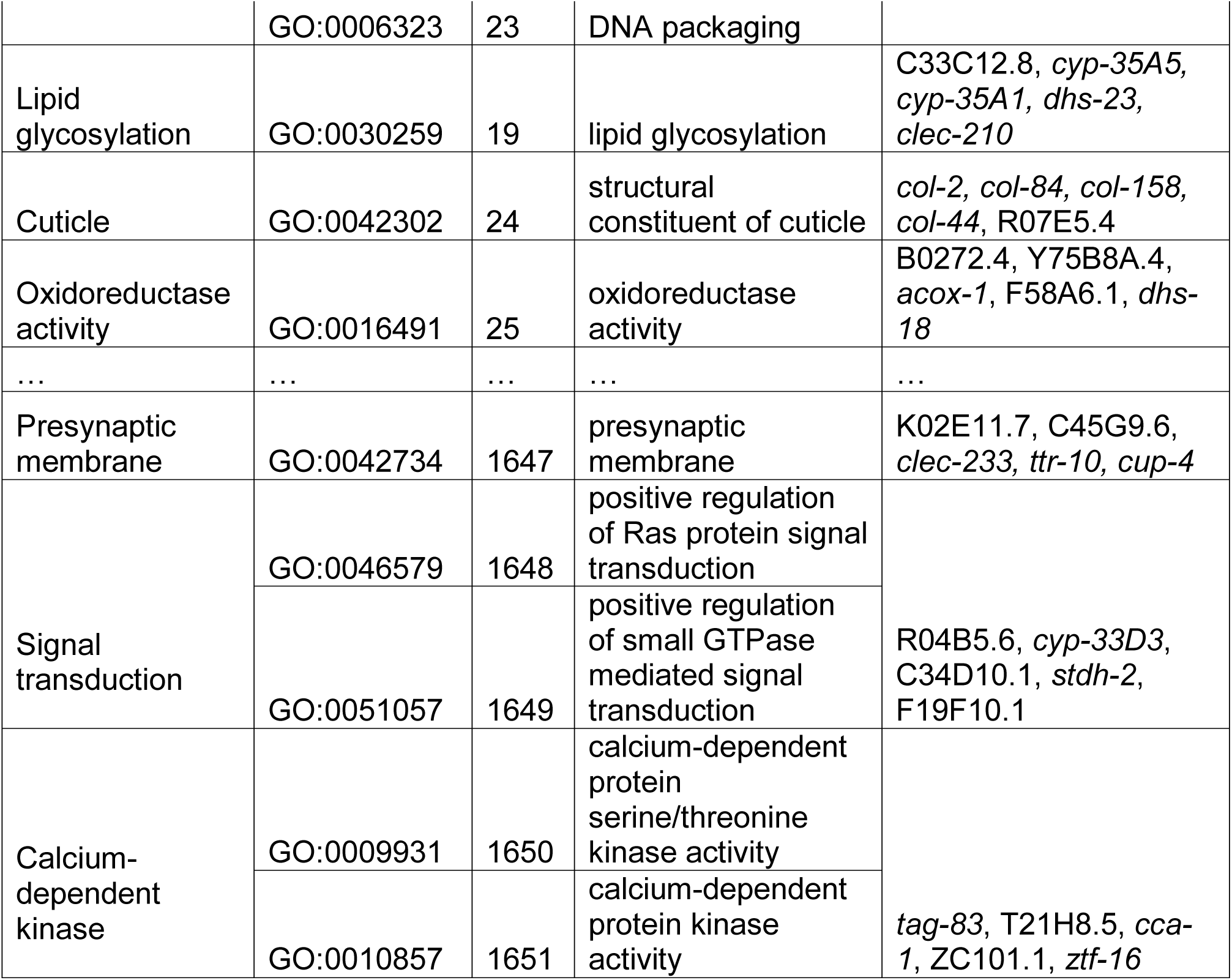
GO categories with strongest and weakest gene weights. The table shows the top 25 and the bottom 5 GO categories, ranked in decreasing order of the weight of their most strongly associated gene prior to normalization. Similar GO categories are grouped together, and the genes with the highest weights for each group are also shown. *rpl-* (<u>r</u>ibosomal <u>p</u>rotein, <u>l</u>arge subunit) and *rps-* (<u>r</u>ibosomal <u>p</u>rotein, <u>s</u>mall subunit) genes encode ribosomal proteins, and *his-* (histone) genes encode nucleosome components.

A sensitive test will tend to give similar results despite small amounts of random noise being added to the input data. To test the sensitivity of a module-weighted annotation-based enrichment test, we obtained gene fold changes for the 5 most recent *C. elegans* Affymetrix experiments deposited to the GEO database. These experiments were not included in the data used to construct the modules. We then added varying amounts of Gaussian noise to the fold changes and for each level of noise, we calculated enrichment z-scores for each GO term using three different methods: the Kolmogorov-Smirnov (KS) test, the t-test (*n.b.*, one-sample and two-sample t-tests produced nearly identical results, data not shown), and scalar projection of the gene fold changes onto a module-weighted GO term matrix. We then compared these results to those obtained by each method when no noise was added (Figure 4a.) Dissimilarity to the initial results (zero added noise) increased rapidly with added noise for both the KS test and t-test, while the projection-based results were similar (ρ > 0.75) at even the highest noise levels tested (5 standard deviations), suggesting that the projection-based test is more sensitive than both the KS and t-tests.

**Figure 4.**
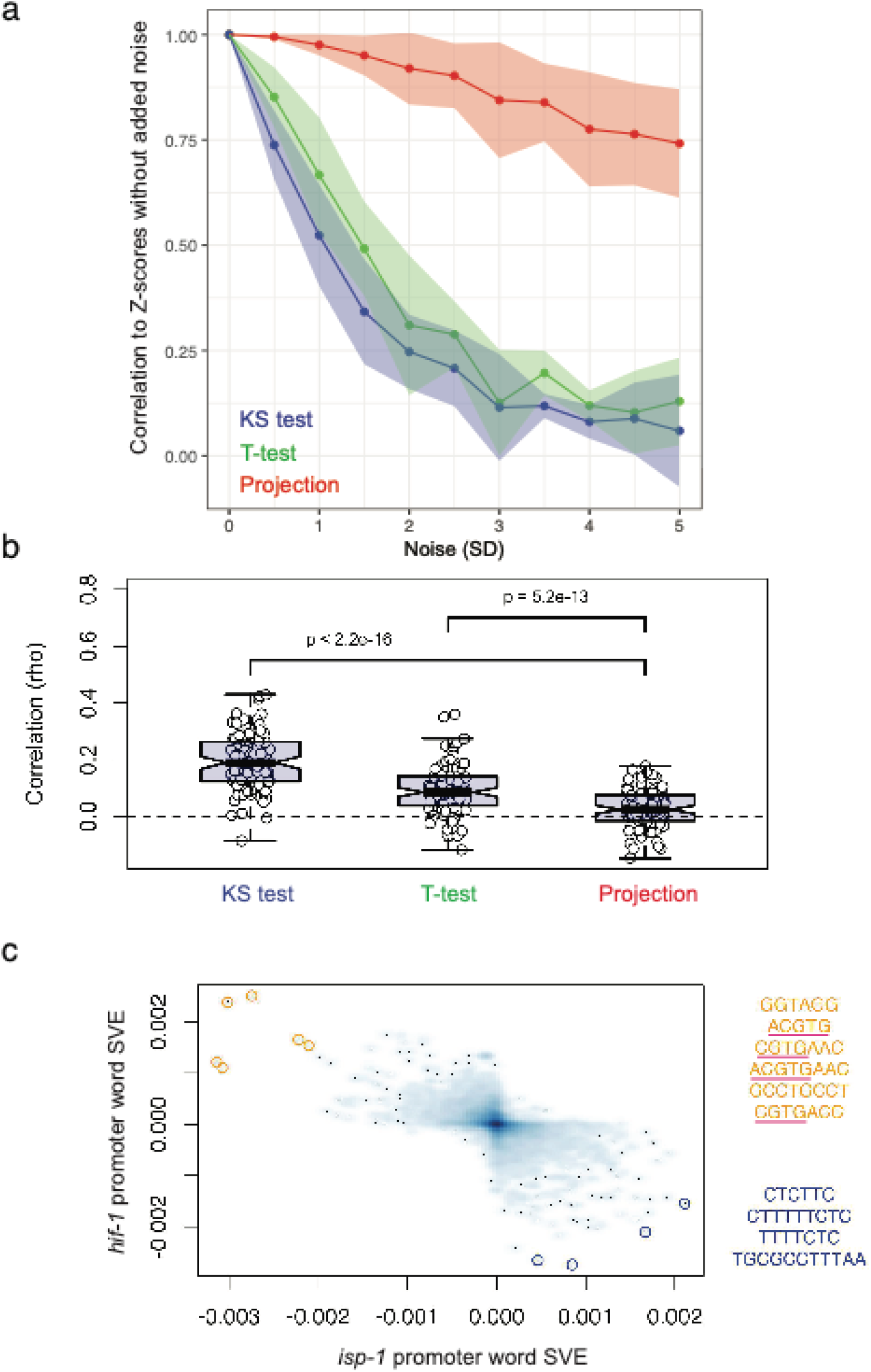
Performance of module-weighted annotation-based tests. (**A**) To assess sensitivity, Gaussian noise was added to gene expression data from 5 recent *C. elegans* Affymetrix experiments not included in the compendium used to train the modules. [The standard deviation of the noise distribution (μ = 0) was varied from 0.5x to 5x the standard deviation of the gene fold change distribution.] At each noise level, z-scores for each GO term (based on 100 random permutations of the gene IDs in the input data) were calculated using three different methods: KS – Kolmogorov-Smirnov test; t-test – two-sample t-test (one-sample test gave highly similar results, data not shown); projection – projection of the gene fold changes into the space defined by the module-weighted annotation matrix, ***R.*** When conducting KS and t-test, fold changes of genes assigned to each GO term were compared to fold changes of genes not assigned to that term. Spearman correlation coefficients were calculated between Z-scores at each noise level and Z-scores without any noise added. (**B**) To assess specificity, 100 highly dissimilar pairs of experiments (Spearman correlation, ρ, of gene fold changes near 0) were selected from a set of 188,805 pairs generated using published microarray data. For each contrast belonging to a pair, we determined Z-scores for GO annotations using the three methods and calculated the rank correlation (rho) of the absolute value of these Z-scores between pair members. The center of the box represents the median value and whiskers extend to the most extreme data point that is not further than 1.5 times the IQR from the box. (**C**) Gene fold changes in *isp-1* and *hif-1* mutants were projection into promoter word space using module-weighted promoter words (see Methods). The 10 points furthest from the origin are highlighted with colored circles in the figure (orange = positive SVE for *hif-1*, blue = negative SVE for *hif-1*), and their words corresponding to these points are shown to the right of the figure. Four of the six words highlighted in orange contain full or partial matches to the canonical *hif-1* binding site, (A/G)CGTG (underlined).

A specific test will tend to show dissimilar results when given dissimilar inputs. To test the specificity of module-weighted annotation analysis, we selected the 100 most dissimilar experiments (correlation of gene fold changes near zero) from a set of 188,805 comparisons. We compared the results of projecting the gene fold changes from each experiment onto the module-weighted annotation matrix to the results obtained from the KS and t-tests and found that the experiments showed the weakest similarity in the significance levels of annotations when using the projection method (p = 5.2e-13, Figure 4b). These results show that annotation enrichment analysis using module-weighted annotations may provide more reliable biological insights than gene set enrichment analyses that rely on the KS (GSEA^41^) or t-tests (PAGE^44^, GAGE^45^).

Module-weighted GO terms for each gene are provided in Supplementary Table 4 and their significance in a query gene expression dataset can be tested using our *C. elegans* gene-modules analysis tools (see Data availability Tool 6).

#### Module-weighted promoter words

While any Boolean gene annotation may be converted into a module-weighted annotation, module-based weights seem particularly well suited for describing regulatory sequences, such as putative transcription factor and microRNA binding sites. To this end, we generated weighted annotations for each of the 5230 words in the promoter word dictionary we constructed using the Mobydick algorithm (described above and in Methods).

To validate predicted regulatory word weights, we searched the literature, the JASPAR database of transcription factor binding profiles^46^, and the GEO database for *C. elegans* transcription factors with both an experimentally-characterized DNA-binding profile and, separately, a microarray experiment that measured gene expression in a loss-of-function mutant of the transcription factor gene. This search yielded 6 transcription factors: *daf-12, daf-16, hif-1, hlh-30, lin-14*, and *nhr-23*. We then projected the loss-of-function microarray experiment data (positively- and negatively-changing genes separately) onto the module-weighted word matrix, ***R*** (Figure 1c), and calculated z-scores for each word. Finally, we compared the top-scoring words to the DNA-binding profiles of the respective transcription factors. If the predicted promoter-word weights are accurate, then words that resemble the binding profile should score highly in this analysis.

For *hif-1* and *nhr-23*, the most significantly enriched words in the positively and negatively changing genes, respectively, matched the canonical binding sites (Table 2). A word matching the *hlh-30* binding site scored 6^th^ overall among the up-regulated genes, and for *daf-12*, four of the top 20 words for the up-regulated genes contained GAACT or AACTT, which partially matched the reverse compliment of a reported *daf-12* binding half-site, AGTTCA^47^. In the *daf-16* data set, several words matching the so-called “*daf-16* associated element” (DAE)^48^ scored highly. However, none of the four words matching the canonical *daf-16* binding site, T(G/A)TTTAC were among the words comprising our Mobydick promoter-word dictionary, precluding these from being represented in the analysis. The canonical binding site for the final transcription factor, *lin-14*, is GAAC, but like the canonical *daf-16* binding site, neither this word nor its reverse compliment was present in the promoter word dictionary, precluding it from representation. Taken together, these results suggest that module weighted regulatory sequences can be used to determine important regulatory sites in a gene expression experiment, and further validate our method for calculating weighted annotations.

**Table 2.**
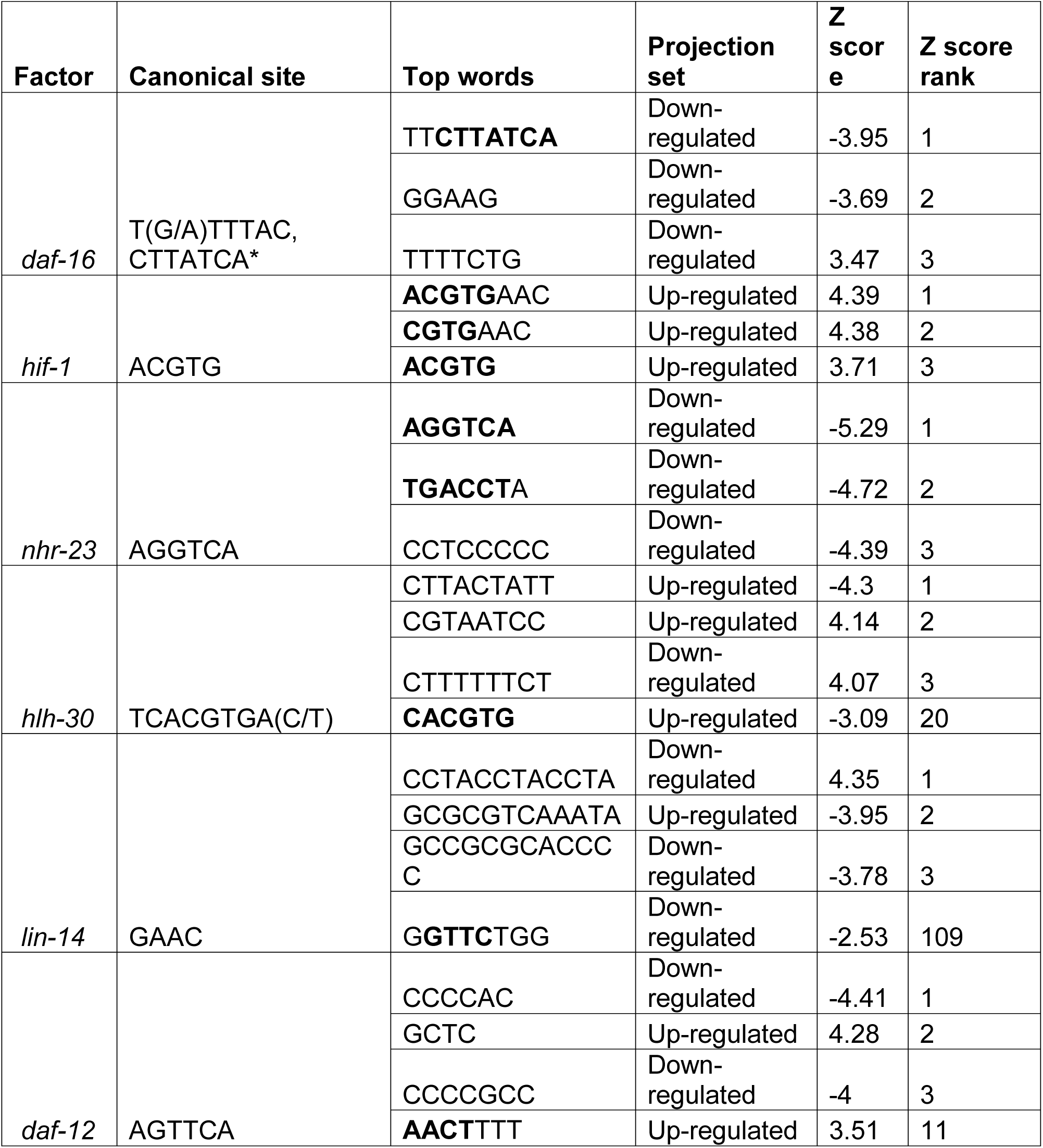
Top scoring promoter words for transcription factor perturbation experiments. The table shows the top three most significant words (those with the most extreme z-scores), calculated using module-weighted promoter words, for each transcription factor loss-of-function microarray experiment. If a full or partial match (4 or more bases) to the canonical binding site of the factor does not occur within the top 3 ranking words, the most significant such match is also shown. Bold letters indicate matching positions to canonical binding sites.

Finally, we tested whether analysis of module-weighted promoter words could reveal the activity of the HIF-1 transcription factor in *isp-1* respiration mutants, missed by analysis of significant genes but implied by analysis of gene module activity (described above). We calculated promoter word z-scores for the *isp-1* microarray data set and compared the results to those for *hif-1*. As predicted, we observed a very strong anti-correlation between the promoter word z-scores for *isp-1* mutants and those for *hif-1* mutants (Figure 4c, R = −0.581, p < 2.2E-16) and four of the six most active words matched the canonical HIF-1 binding site [(A/G)CGTG; underlined in Figure 4c]. These results further support the utility of module-weighted promoter oligonucleotides for identifying biologically relevant regulatory sequences directly from gene expression data.

#### Module-weighted annotations as a hypothesis generation tool

Our process of weighting gene annotations using modules provisionally transfers annotations that are enriched in a module to all gene members of that module (proportional to each gene’s inclusion). This can be useful for studying individual genes (see Data availability Tool 5 and Supplementary Table 4) – if a query gene has a strongly-weighted association to a GO term with which it is not traditionally associated, it means that this gene is co-expressed with other genes that *are* traditionally associated with the GO term. For example, most of the module-weighed GO terms associated with the small ribosomal subunit S16 (*rps-16)* have something to do with the ribosome or translation (see Data availability Tool 5), implying that *rps-16* is tightly co-expressed with other genes involved in ribosome biogenesis and not much else. On the other hand, while *sod-3*, a superoxide dismutase, does have significant weights for GO terms with which it is traditionally associated (e.g. oxidoreductase activity, superoxide metabolic process and response to superoxide), catalase activity ranks more highly for *sod-3*. This means that *sod-3* is co-expressed with genes annotated as catalases. While it is unlikely that *sod-3* has a novel catalase activity, it is more likely that expression of *sod-3*, a superoxide dismutase, is coordinated with expression of catalases, since hydrogen peroxide produced by *sod-3* is further degraded by catalases to avoid damage to the cell. Some of the other top-ranking module-weighted GO terms for *sod-3* describe dioxygenase activity. Similarly, this may indicate that dioxygenases generate superoxide radicals, and therefore increased expression of these enzymes is typically correlated with increased expression of a superoxide dismutase. Therefore, analysis of module-weighted annotations of individual genes can help form hypotheses about novel gene functions and transcriptional co-regulation of distinct biological processes.

## Discussion

We have captured much of the transcriptional wiring of *C. elegans* by extracting gene co-expression modules from a large compendium of data. These 209 modules represent transcriptional signatures of diverse biological processes that can occur in *C. elegans* and, along with the tools we provide, can be used as a resource for analyzing gene expression data. Because genes are grouped into modules based purely on how they behave in experimental assays, and because modules encompass both previously-annotated and unannotated genes alike, modules can reveal signals within transcriptomic data that otherwise would be missed due to incomplete knowledge of gene function, subtle gene expression changes or noisy data. Experimentally, gene-expression modules can be used to deconvolve complex phenomena into subsets of co-regulated genes, genes that likely act together to mediate a specific process. The modules can be annotated extensively, as we have done, and these annotations can be applied provisionally to all genes in the module. Regulatory factors associated with specific annotations (like 5’ or 3’ oligonucleotide words) can be implicated as well, and functional links can be revealed between dissimilar conditions that activate the same modules.

ICA has been applied to the prediction of gene modules before, but we could find no examples in the literature of optimizing the process using biological metrics in the manner that we describe. Combined with the improved ability to partition independent components provided by our artificial neural network approach, we expect that DEXICA will be useful for constructing gene modules for other organisms. The DEXICA software and our *C. elegans* modules and data are freely available as R packages online (see Data availability).

While we have taken steps to maximize module prediction accuracy for the microarray compendium we assembled, many additional gene modules may exist in *C. elegans* that were not perturbed sufficiently in the samples comprising the compendium to be detected. These gene modules would remain hidden. As new areas of research are explored and new experiments are published, however, new fundamental gene modules may be discovered. For example, most of the gene expression data in our compendium was collected using whole animals. Data collected from isolated cells or tissues could help produce modules that are active in a relatively small number of cells. The DEXICA package can be used to create new and improved gene modules using compendia with expanded information content.

The impetus for constructing gene modules in *C. elegans* was to create gene groupings that do not rely on the existing annotations, because annotations of many genes are missing or incomplete. We then wondered whether numeric scores based on gene co-expression could actually improve the existing gene annotations, leading us to develop the concept of module-weighted annotations. By weighting an association between each gene and each annotation by the degree to which that annotation appears to predict gene modularity, annotations that are shared by co-expressed genes are “boosted” and those that are not are diminished. It is important to note, however, that relevance scores between genes and annotations could be calculated using other metrics of gene behavior as well (e.g. a gene might be weakly associated with a term in the context of gene expression but strongly associated with it in the context of physical protein interactions).

We have found that analysis of module-weighted GO terms is less sensitive to noise than are typical statistical over-representation tests. Furthermore, because module-weighted annotations incorporate information about gene expression, they effectively model the gene-gene correlation structure of the system. This is useful because typical over-representation tests do not perform well with gene sets that have a high level of gene-gene correlation (i.e. annotations assigned to genes that are strongly co-expressed are more likely to be significant^49–51^. GSEA, for example, deals with gene-gene correlation issue using permutation. We show that module-weighted GO terms produce significantly fewer false positives than do over-representation tests (see Figure 4b). Therefore, we think that module-weighted annotations are a promising new way to address the gene-gene correlation problem and are working to develop it further for annotation enrichment analysis.

Module-weighted annotations may be especially useful for promoter/3’-UTR word analysis. Projection of gene expression changes onto the module-weighted word matrix allows identification of potential regulatory sequences directly from a list of gene fold changes, bypassing many steps required by traditional regulatory sequence detection (sequence retrieval, repeat masking, over-representation analysis, and background correction). More importantly, this analysis is more likely to yield true positive results because it makes use of a large amount of historical data (via the modules) to filter out sequences with no apparent regulatory function.

We used module-weighted word analysis to analyze data from six transcription factor perturbation microarray experiments. Four of the six produced high scoring promoter words that closely matched the known DNA binding sites of the corresponding transcription factors. For the two that did not, *daf-16* and *lin-14*, words that exactly matched the factors’ canonical binding sites were not present in the promoter word dictionary generated by the Mobydick algorithm, and results for these factors could be poor for this reason. Interestingly, the second highest scoring word for the genes down-regulated upon *daf-16* perturbation was GGAAG, and this word occurs twice more as a substring among the top 20 scoring words. This sequence is a partial match to an alternative *daf-16* binding site reported in hookworm^52^, G(A/G)(C/G)A(A/T)G, suggesting that this site may be functional in *C. elegans* as well.

Finally, because annotations shared among a significant fraction of the genes in a module are “transferred” to all genes in the module, module-weighted annotations can be used provisionally to infer novel functions for genes, which is especially useful for studying poorly annotated genes.

## Methods

### Compendium construction

To build the compendium of 1386 *C. elegans* Affymetrix arrays, we first downloaded all CEL files with the appropriate platform ID (GPL200) from the GEO database available on March 1, 2014, excluding those for which the organism was not *C. elegans* and the sample type was not RNA. We excluded arrays from experiments for which fewer than 8 hybridizations were performed in order to mitigate the effect that under-sampled conditions might have on predicted modules. We then performed a quality control step using the quality assessment functions provided in the *simpleAffy* (v2.40.0) R package (http://bioinformatics.picr.man.ac.uk/simpleaffy/), discarding arrays that did not meet the quality thresholds recommended in the *simpleAffy* documentation.

We generated expression values for probesets separately for each experiment (determined by GEO series IDs) using the RMA preprocessing procedure provided in the *affy* (v1.40.0) R package^53^, then used the *bias* (v0.0.5) R package^54^ to remove intensity-dependent biases in expression levels. We then concatenated the expression matrices for each experiment into a single matrix. Next, we either performed between-experiment quantile normalization on the entire matrix using the *limma* (v3.18.13) R package^55^, or omitted this step, depending on preprocessing method to be tested. Finally, we scaled and centered the arrays and centered the genes such that the mean of each row and column were zero and the standard deviation of each array was 1.

### Conducting ICA

To conduct ICA of the gene expression matrix, we used the *fastICA* (v1.2-0) R package (http://CRAN.R-project.org/package=fastICA) with default parameters except for the “method” parameter, which we set to “C” to increase computational speed, and the “row.norm” parameter, which we set to “TRUE” in order to balance the total compendium variance between genes with subtle changes in expression values and those with large changes in expression values.

### Partitioning of independent components

To convert independent components to discrete sets of genes, we employed two methods. In the first, for each component, we assigned all genes with a weight <= −3 to the negative hemi-module, and all genes with a weight >= 3 to the positive hemi-module. In the second, we created an artificial neural network using the *neuralnet* (v1.32) R package (http://CRAN.R-project.org/package=neuralnet) to predict positive and negative partitioning thresholds for each independent component, based on the component’s skewness and kurtosis (see Supplementary Methods), then assigned genes whose weights exceeded these thresholds to the corresponding hemi-modules.

### Obtaining gene annotations and microarray data

To obtain GO term and REACTOME pathway annotations for genes we used the *biomaRt* (v2.18.0) R package^56^, using the *ensembl* mart for data retrieval. To obtain tissue annotations for *C. elegans* genes, we downloaded all available data from the GFP Worm database (http://gfpweb.aecom.yu.edu/)^28^, which contains annotated expression patterns of promoter::GFP fusion constructs; in total, this dataset provided annotations for 1821 genes across 89 tissue types (*n.b.*, we considered the same tissue in different development stages to be distinct tissue types). To obtain fold changes for *isp-1* mutants, we used data previously published by our group in which *isp-1(qm150)* mutants were compared to wild type controls^57^. To obtain fold changes for *hif-1* mutants, we used the *maanova* (v1.33.2) R package (http://research.jax.org/faculty/churchill) and data previously published by Shen, et al.^58^, to calculate the induced gene fold changes upon mutation of *hif-1*. All other microarray data was obtained directly from the authors of the original publications of from the GEO database (see Supplementary Methods).

### Optimizing gene module prediction

To optimize gene module prediction, we performed ICA with a variety of different data preprocessing options (e.g., the choice of preprocessing algorithm (RMA, GCRMA, PLIER, MAS 5.0), background, perfect match, bias correction, and normalization methods), and with a varied number of extracted components from 5 to 500 by increments of 5. For each parameter combination, we repeated ICA 5 times.

We tested the biological validity of the independent components generated by each ICA run by determining the number of annotations that were enriched in at least one hemi-module. We chose not to optimize based on the number of modules with at least one significant annotation, as tests using simulated data showed that this approach could lead to signals being split into multiple, less accurate representations (data not shown). We first calculated a p-value for the enrichment of genes associated with each annotation term in each hemi-module using the hypergeometric test. We then applied the Simes method^59^ for multiple hypothesis testing (alpha = 0.05) to the set of p-values for each annotation term. The Simes method is similar to the Benjamini-Hochberg method^60^ for controlling the false discovery rate, but differs in a way that makes it more appropriate here: it aims to answer the question, “Given a set of p-values, what is the likelihood that at least one null hypothesis is false?”, while the Benjamini-Hochberg method asks, “What fraction of rejected null hypotheses are actually true (i.e., falsely discovered)?” Failure of the Simes test indicates that at least one null hypothesis is false at the specified alpha level. To verify the accuracy of our module quality statistics, we repeated all tests using module definition matrices in which gene IDs had been randomly shuffled.

### Quantification of module activity in gene expression data

To project a data vector, **x**, such as a set of gene expression fold changes, onto a set of gene modules, we used the scalar projection method, in which a mixing vector, **a**, is calculated from the dot product of the data vector and the unit vectors comprising the module definitions, 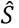, as shown in equation 2:

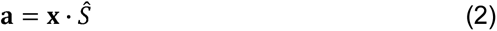

The resulting mixing vector, ***a***, provides an indication of the weight of each module definition vector in the projected data, **x**. Projection of a data matrix, ***X***, which generates a mixing matrix, ***A***, was carried out using the same procedure. To calculate signed variance explained (SVE), we calculated the relative variance explained (VE) for each module from ***a*** as follows, where *n* is the total number of modules:

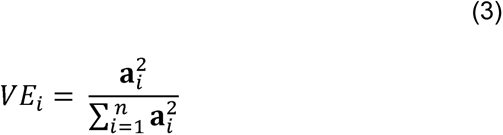

We then multiplied these values, which are strictly positive, by −1 in each case where **a_i_** < 0 to obtain SVE.

### Generation of *E* matrix

First, ANN-based partitioning of the module definition matrix, ***S***, was used to generate matrix ***S_p_***. ***S_p_*** contains two gene sets per independent component (which we refer to as hemi-modules), for a total of 418. Using a matrix of known Boolean associations between genes and annotations, ***B***, we calculated a hypergeometric probability for each annotation in each hemi-module. We used the frequency of genes bearing a particular annotation in the hemi-module, the frequency of such genes in the compendium, the number of genes in the hemi-module, and the number of genes not in the hemi-module as the *q, m, k*, and *n* input parameters, respectively, to the *phyper()* function of the *stats* (v3.0.3) R package (http://www.R-project.org/).

We used these p-values to populate a matrix, ***E***, with a row for each hemi-module and a column for each annotation. For under-represented annotations, we entered the *log(p-value)* in the matrix, and for over-represented annotations we entered the *–log(p-value)*.

### Generation of module-weighted annotations

Our calculation of module-weighted annotations takes advantage of the fact that, in modules generated by ICA, prior to partitioning, each gene has a weight in each module. Given a score or weight for each annotation in each module, this allows genes to be associated with annotations via a matrix product calculation. In the ***E*** matrix (see above) highly positive values correspond to strongly enriched annotations in a hemi-module, highly negative values correspond to strongly depleted annotations. We transform the gene module matrix, ***S***, into an unpartitioned hemi-module matrix, ***H***, by concatenating it with a negative copy of itself column-wise:

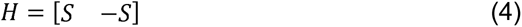

The product of this matrix with the ***E*** matrix produces matrix ***R***, which relates genes to annotations:

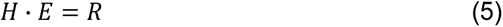

As a final step, we normalize the values in this matrix row-wise, i.e, separately for each annotation, by subtracting the mean and dividing by the standard deviation.

### Analysis of GO terms based on gene weights

To test whether the ranking of GO terms based on gene weights (Table 1) is statistically significant, we constructed a list comprising all words used in all GO terms, excluding words shorter than three letters and uninformative words (e.g. “the” and “for”.) We then tested each word for bias toward appearing near the top or bottom of the ranked GO term list. In agreement with our initial observations, the most significantly top-biased words pertained to macromolecular complexes, such as “nucleosome”, “cilium”, and “ribosomal”, and the most significant bottom-biased words pertained to cell signaling, such as “signal”, “kinase”, and “receptor”. Many of the GO terms containing cell signaling words were generic in nature, e.g., protein kinase regulator activity, thus, our results may partially be explained by a lack of co-regulation among constituents of different signaling pathways. However, some specific cell signaling terms, e.g., *Notch signaling pathway* also appeared near the bottom of the ranked GO term list, suggesting that the genes annotated with such terms are either not strongly co-regulated at the gene expression level or that the biological conditions represented by the compendium did not perturb their expression enough to form modules with our method.

### Analysis of expression data with module-weighted annotations

To test whether a set of gene fold-changes were significantly enriched for specific annotations, given a module-weighted annotation matrix, ***R***, we calculated the dot product of the data vector, **x**, comprising the set of gene fold-changes, and the ***R*** matrix:

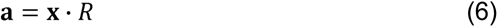

The resulting vector, **a**, provides an indication of the degree to which genes with strong weights for each annotation also have strong fold-changes. To generate p-values from these, we permuted the fold-change vector, ***x***, 1000 times to create a background distribution for each annotation, which we then used to determine z-scores.

### Data availability

The authors declare that all data supporting the findings of this study are available within the paper and its supplementary tables. Dynamic access to the data is also provided through a graphical user interface at http://genemodules.org/. Specifically, 6 separate functionalities are provided:

**Tool 1**: Determine which of the 209 modules defined here are active in the user’s gene expression data (by inputting a list of gene-expression fold changes)

**Tool 2**: Using the partitioned 209 modules as gene sets, test whether a list of user-specified genes is enriched in any particular module

**Tool 3**: Detailed description of each of the 209 *C. elegans* gene modules

**Tool 4**: Visualize how genes assigned to a module change under a variety of conditions (conditions can be chosen from the 716 perturbations derived from our microarray compendium or provided by a user)

**Tool 5**: Get module-weighted GO terms associated with a user-specified gene

**Tool 6:** Identify the top module-weighted GO terms in the user’s gene expression data

### Code availability

Code described in this study is available as an R package: DEXICA R package https://github.com/MPCary/DEXICA. A small dataset to demo the package can be downloaded from https://github.com/MPCary/DEXDATA.Celegans.

~~~
To install:
install.packages(“devtools”)
library(devtools)
install_github(“MPCary/DEXICA”, build_vignettes = TRUE)
install_github(“MPCary/DEXDATA.Celegans”)
~~~

## Acknowledgements

We would like to thank H. Li, H. El-Samad, H. Madhani and Frank Li for critical review and discussion of this work. This work was supported by the NIH R36/R01 grant AG011816 to C. K.

## Author contributions statement

M.C., K.P. and C.K. conceived the study and wrote the manuscript. M.C. and K.P. performed data analyses. M.C. developed the R package. K.P. developed the web-based tools. C.K. secured funding.

## References

1. Lee, R. Y. N. et al. WormBase 2017: molting into a new stage. Nucleic acids research 46, D869–D874

2. Ashburner, M. et al. Gene ontology: tool for the unification of biology. The Gene Ontology Consortium. Nature genetics 25, 25–29 (2000).

3. Kanehisa, M., Goto, S., Sato, Y., Furumichi, M. & Tanabe, M. KEGG for integration and interpretation of large-scale molecular data sets. Nucleic acids research 40, D109–14

4. Fabregat, A. et al. The Reactome Pathway Knowledgebase. Nucleic acids research 46, D649–D655

5. Wu, W.-S., Li, W.-H. & Chen, B.-S. Computational reconstruction of transcriptional regulatory modules of the yeast cell cycle. BMC Bioinformatics 7, 421 (2006).

6. Wang, W. et al. Inference of combinatorial regulation in yeast transcriptional networks: a case study of sporulation. PNAS 102, 1998–2003 (2005).

7. Teschendorff, A. E., Journée, M., Absil, P. A., Sepulchre, R. & Caldas, C. Elucidating the altered transcriptional programs in breast cancer using independent component analysis. PLoS Comput. Biol. 3, e161 (2007).

8. Mostafavi, S. et al. A molecular network of the aging human brain provides insights into the pathology and cognitive decline of Alzheimer’s disease. Nature Neuroscience 21, 811–819 (2018).

9. Hughes, T. R. et al. Functional Discovery via a Compendium of Expression Profiles. Cell 102, 109–126 (2000).

10. Engreitz, J. M., Daigle, B. J., Marshall, J. J. & Altman, R. B. Independent component analysis: mining microarray data for fundamental human gene expression modules. J Biomed Inform 43, 932–944 (2010).

11. Zhou, W. & Altman, R. B. Data-driven human transcriptomic modules determined by independent component analysis. BMC Bioinformatics 19, 327 (2018).

12. Kim, S. K. et al. A Gene Expression Map for Caenorhabditis elegans. Science 293, 2087–2092 (2001).

13. Gerstein, M. B. et al. Architecture of the human regulatory network derived from ENCODE data. Nature 489, 91–100 (2012).

14. Oeckinghaus, A., Hayden, M. S. & Ghosh, S. Crosstalk in NF-κB signaling pathways. Nat. Immunol. 12, 695–708 (2011).

15. Lee, S.-I. & Batzoglou, S. Application of independent component analysis to microarrays. Genome Biol. 4, R76 (2003).

16. Gong, T. et al. Gene module identification from microarray data using nonnegative independent component analysis. Gene Regul Syst Bio 1, 349–363 (2007).

17. Rustici, G. et al. ArrayExpress update--trends in database growth and links to data analysis tools. Nucleic acids research 41, D987–90

18. Barrett, T. et al. NCBI GEO: archive for functional genomics data sets--10 years on. Nucleic acids research 39, D1005–10 (2011).

19. Ihmels, J., Bergmann, S. & Barkai, N. Defining transcription modules using large-scale gene expression data. Bioinformatics 20, 1993–2003 (2004).

20. Segal, E., Yelensky, R. & Koller, D. Genome-wide discovery of transcriptional modules from DNA sequence and gene expression. Bioinformatics 19 Suppl 1, i273–82 (2003).

21. Michoel, T., De Smet, R., Joshi, A., Marchal, K. & Van de Peer, Y. Reverse-engineering transcriptional modules from gene expression data. Ann N Y Acad Sci 1158, 36–43 (2009).

22. Saelens, W., Cannoodt, R. & Saeys, Y. A comprehensive evaluation of module detection methods for gene expression data. Nature communications 9, 1090 (2018).

23. Teschendorff, A. E., Journee, M., Absil, P. A., Sepulchre, R. & Caldas, C. Elucidating the altered transcriptional programs in breast cancer using independent component analysis. PLoS Comput. Biol. 3, e161 (2007).

24. Zhou, W. & Altman, R. B. Data-driven human transcriptomic modules determined by independent component analysis. BMC Bioinformatics 19, 327 (2018).

25. Purdom, E. & Holmes, S. P. Error distribution for gene expression data. Stat Appl Genet Mol Biol 4, Article16 (2005).

26. Hyvärinen, A. & Oja, E. Independent Component Analysis: Algorithms and Application. Neural Networks 13, 411–430 (2000).

27. Kong, W., Vanderburg, C. R., Gunshin, H., Rogers, J. T. & Huang, X. A review of independent component analysis application to microarray gene expression data. Biotechniques 45, 501–520 (2008).

28. Hunt-Newbury, R. et al. High-throughput in vivo analysis of gene expression in Caenorhabditis elegans. PLoS biology 5, e237 (2007).

29. Lee, S.-I. & Batzoglou, S. Application of independent component analysis to microarrays. Genome Biol. 4, R76 (2003).

30. Chiappetta, P., Roubaud, M. C. & Torresani, B. Blind source separation and the analysis of microarray data. J Comput Biol 11, 1090–1109 (2004).

31. Frigyesi, A., Veerla, S., Lindgren, D. & Hoglund, M. Independent component analysis reveals new and biologically significant structures in micro array data. BMC Bioinformatics 7, 290 (2006).

32. Bussemaker, H. J., Li, H. & Siggia, E. D. Building a dictionary for genomes: identification of presumptive regulatory sites by statistical analysis. PNAS 97, 10096–10100 (2000).

33. Zaslaver, A., Baugh, L. R. & Sternberg, P. W. Metazoan operons accelerate recovery from growth-arrested states. Cell 145, 981–992

34. Mayr, C. & Bartel, D. P. Widespread shortening of 3’UTRs by alternative cleavage and polyadenylation activates oncogenes in cancer cells. Cell 138, 673–684 (2009).

35. Baugh, L. R. To grow or not to grow: nutritional control of development during Caenorhabditis elegans L1 arrest. Genetics 194, 539–555

36. Dancy, B. M., Sedensky, M. M. & Morgan, P. G. Effects of the mitochondrial respiratory chain on longevity in C. elegans. Exp Gerontol 56, 245–255

37. Baker, B. M., Nargund, A. M., Sun, T. & Haynes, C. M. Protective coupling of mitochondrial function and protein synthesis via the eIF2alpha kinase GCN-2. PLoS genetics 8, e1002760

38. Durieux, J., Wolff, S. & Dillin, A. The cell-non-autonomous nature of electron transport chain-mediated longevity. Cell 144, 79–91

39. Pellegrino, M. W. et al. Mitochondrial UPR-regulated innate immunity provides resistance to pathogen infection. Nature 516, 414–417

40. Nargund, A. M., Pellegrino, M. W., Fiorese, C. J., Baker, B. M. & Haynes, C. M. Mitochondrial import efficiency of ATFS-1 regulates mitochondrial UPR activation. Science 337, 587–590

41. Subramanian, A., et al. Gene set enrichment analysis: a knowledge-based approach for interpreting genome-wide expression profiles. PNAS 102, 15545–15550 (2005).

42. Lee, S. J., Hwang, A. B. & Kenyon, C. Inhibition of respiration extends C. elegans life span via reactive oxygen species that increase HIF-1 activity. Current biology: CB 20, 2131–2136

43. Wade, C. H., Umbarger, M. A. & McAlear, M. A. The budding yeast rRNA and ribosome biosynthesis (RRB) regulon contains over 200 genes. Yeast 23, 293–306 (2006).

44. Kim, S. Y. & Volsky, D. J. PAGE: parametric analysis of gene set enrichment. BMC Bioinformatics 6, 144 (2005).

45. Luo, W., Friedman, M. S., Shedden, K., Hankenson, K. D. & Woolf, P. J. GAGE: generally applicable gene set enrichment for pathway analysis. BMC Bioinformatics 10, 161 (2009).

46. Mathelier, A. et al. JASPAR 2016: a major expansion and update of the open-access database of transcription factor binding profiles. Nucleic acids research

47. Shostak, Y., Van Gilst, M. R., Antebi, A. & Yamamoto, K. R. Identification of C. elegans DAF-12-binding sites, response elements, and target genes. Genes & development 18, 2529–2544 (2004).

48. Zhang, P., Judy, M., Lee, S. J. & Kenyon, C. Direct and indirect gene regulation by a life-extending FOXO protein in C. elegans: roles for GATA factors and lipid gene regulators. Cell metabolism 17, 85–100 (2013).

49. Goeman, J. J. & Buhlmann, P. Analyzing gene expression data in terms of gene sets: methodological issues. Bioinformatics 23, 980–987 (2007).

50. Gatti, D. M., Barry, W. T., Nobel, A. B., Rusyn, I. & Wright, F. A. Heading down the wrong pathway: on the influence of correlation within gene sets. BMC Genomics 11, 574 (2010).

51. Tamayo, P., Steinhardt, G., Liberzon, A. & Mesirov, J. P. The limitations of simple gene set enrichment analysis assuming gene independence. Stat Methods Med Res 25, 472–487 (2016).

52. Gao, X. et al. Identification of hookworm DAF-16/FOXO response elements and direct gene targets. PloS one 5, e12289

53. Gautier, L., Cope, L., Bolstad, B. M. & Irizarry, R. A. affy--analysis of Affymetrix GeneChip data at the probe level. Bioinformatics 20, 307–315 (2004).

54. Eklund, A. C. & Szallasi, Z. Correction of technical bias in clinical microarray data improves concordance with known biological information. Genome Biol. 9, R26 (2008).

55. Smyth, G. K. in Bioinformatics and Computational Biology Solutions using R and Bioconductor (eds. Gentleman, R., Carey, V., Dudoit, S., Irizarry, R. & Huber, W.) 397–420 (Springer, 2005).

56. Durinck, S., Spellman, P. T., Birney, E. & Huber, W. Mapping identifiers for the integration of genomic datasets with the R/Bioconductor package biomaRt. Nat Protoc 4, 1184–1191 (2009).

57. Cristina, D., Cary, M., Lunceford, A., Clarke, C. & Kenyon, C. A regulated response to impaired respiration slows behavioral rates and increases lifespan in Caenorhabditis elegans. PLoS genetics 5, e1000450 (2009).

58. Shen, C., Nettleton, D., Jiang, M., Kim, S. K. & Powell-Coffman, J. A. Roles of the HIF-1 hypoxia-inducible factor during hypoxia response in Caenorhabditis elegans. The Journal of biological chemistry 280, 20580–20588 (2005).

59. SImes, R. J. An Improved Bonferroni Procedure for Multiple Tests of Significance. Biometrika 73, 751–754 (1986).

60. Benjamini, Y., B, Y. H. J. O. T. R. S. S. S.1995. Controlling the false discovery rate: a practical and powerful approach to multiple testing. JSTOR doi:10.2307/2346101

