## Supplementary materials for "A resource for analyzing *C. elegans*’ gene expression data using transcriptional gene modules and module-weighted annotations"

**Supplementary discussion**

*Theoretical basis of module-weighted annotations*

Almost invariably in gene databases, annotations are connected to genes by a Boolean relationship, e.g., a GO term either *is* or *is not* associated with a particular gene. However, Boolean annotations cannot provide an indication of the relative strength of the association between an annotation and a gene. For example, *in vitro* transcription factor binding assays, such as those generated by the ENCODE^1^ and modENCODE^2^ projects, can be used to annotate genes with lists of transcription factors that bind proximally. However, the relative importance of each factor to the regulation of each gene would not be clear from such data alone. Supplementing those annotations with numeric values, on the other hand, could be used to provide an indication of how much control a transcription factor appears to exert over the expression of an associated gene.

Gene expression modules provide an indication of how the cell’s gene regulation is “wired” by revealing which genes tend to be co-expressed. This makes it possible to compute a score for each gene-annotation pair from the degree to which genes assigned the common annotation are actually co-expressed.

Another well-documented shortcoming of Boolean annotations arises when researchers ask whether certain annotations (e.g. GO terms) are over-represented among differentially expressed genes. Over-representation analyses typically start with a statistical test, such as a Kolmogorov-Smirnov test (GSEA^3^) or a t-test (PAGE^4^, GAGE^5^), that assumes statistical independence between genes. This is followed by a correction procedure (e.g., by comparing results to a reference distribution generated from random permutations of the input data), made necessary by the fact that expression levels of genes are not, in fact, statistically independent. On the contrary, genes in many GO categories are known to be highly correlated which, if left unaccounted for, can lead to substantial numbers of false positive results^6-8^. This problem can be conceptualized by reflecting on the fact that, due to biological stochasticity, a particular gene might be slightly higher in experimental samples relative to controls. If expression of this gene is tightly correlated with some other genes, they too are more likely to be slightly higher. If all these genes share a common annotation, this annotation is likely to be declared significant. Current methods to correct for this effect depend on having a sufficient number of replicate samples for use in generating a reference distribution, and the degree to which false positives arising from gene-gene correlations survive the correction procedure is generally unclear. In cases where a user only has a list of gene fold changes, rather than sample data for each biological replicate, correcting for the effect of gene-gene correlations becomes more problematic.

We hypothesized that since gene-expression modules capture the complex gene set correlation structure of a system, they could be used to improve the identification of biologically significant annotations from gene expression data. Unlike typical annotation enrichment tests, analysis of gene expression data using module-weighted annotations can bypass the statistical test step entirely (see Methods and Data availability Tool 5), instead starting from a scalar projection of gene fold changes into the space defined by the module-weighted annotation matrix (matrix ***R***, Figure 1c). The resulting values (one for each annotation) would provide an indication of the degree to which genes with strong weights for each annotation also have strong fold-changes. Thus, tightly co-regulated genes that share an annotation would be declared significant only if they also have substantial fold-changes (which is likely to be an indication of biological significance).

**Supplementary methods**

**Generation of simulated data**

To generate a simulated microarray compendium containing a proscribed number, *n*, of true gene modules, we performed ICA on a test compendium and extracted 2*n* independent components. We then randomly selected *n* rows of the resulting S matrix and multiplied these with the corresponding columns of the A matrix. This generated a matrix with the same dimensions as the test compendium, but with only *n* latent gene modules. We then added random Gaussian noise to each data point in this matrix. Increasing the level of noise decreased the accuracy of modules extracted from the matrix, but did not alter the observed property that accuracy reached a maximum when the number of extracted components matched the number of latent modules (data not shown).

**Construction of Mobydick dictionaries**

To construct promoter and 3’-UTR dictionaries, we ran the Mobydick^9^ program once on the complete set of *C. elegans* promoters, using DNA sequence from the transcription start site to 2000 b.p. upstream for each gene, and again on the complete set of 3’-UTRs with lengths of at least 25 n.t. Sequences were obtained using the *biomaRt* (ver 2.14.0) R package^10^. Application of Mobydick to promoter sequences produced a dictionary of 5230 words, and application to 3’-UTR sequences produced a dictionary of 968 words.

**Calculation of significance of 3’-UTR word enrichment**

Because 3’-UTRs differ in length, and because gene modules show a tendency toward inclusion of genes with similar length 3’-UTRs, calculation of the enrichment of 3’-UTR words in module genes required a length-normalization step. To achieve this, we applied the method described van Helden, et al.^11^. Briefly, we determined the per nucleotide frequency of each word in the entire set of 3’-UTRs, then used the binomial distribution to determine whether each word occurs more often than expected by random chance in a sequence, given the number of occurrences of the word in the sequence and the sequence length. We then applied the Holm-Bonferroni correction to the resulting p-values and marked all words with a corrected p-value < 0.5 (i.e., less than a 50% chance of seeing the number of occurrences by random chance) as present in the 3’-UTR.

**Statistical testing of module 3’-UTR length bias**

We observed that *C. elegans* 3’-UTR lengths are approximately log-normally distributed (Supplementary Figure 6). Therefore, we chose to use the log of each 3’-UTR length in our calculations to allow the use of parametric statistical tests, such as Student’s t-test. For those genes with multiple annotated 3’-UTRs, we determined the log of the individual 3’-UTR lengths and used the mean of these numbers for the gene’s 3’-UTR length.

In our statistical test for 3’-UTR length biases in predicted modules, we first calculated the weighted mean *C. elegans* 3’-UTR length. We weighted each gene’s contribution to this mean by the frequency with which it appears in our predicted modules in order to adjust for different propensities for module inclusion by different genes. We then used one-sample t-tests to calculate p-values for whether the mean 3’-UTR length of each hemi-module differs significantly from the weighted mean *C. elegans* 3’-UTR length. We used the Benjamini-Hochberg procedure on these p-values to control the false discovery rate at a level of 0.1.

**Generation of Affymetrix contrasts**

To generate the 716 Affymetrix contrasts, we examined all *C. elegans* Affymetrix experiments (organized as GEO series) available for download in the GEO database on January 23, 2015. For each experiment, we generated contrasts (two sets of microarrays to compare to each other) based on the following rules: 1) we compared genetic mutants to wild type controls, or to the genotype most resembling wild type if true wild type animals were not used in the experiment, 2) we compared animals subjected to a treatment, such as an environmental stress, to animals with the same genotype but subjected to control conditions, 3) we compared animals harvested at different time points to animals with the same genotype but harvested at the earliest time point represented in the experiment. When necessary, we divided large experiments into smaller subsets in order to simplify contrast generation and facilitate the application of the above rules. We generated fold-changes from contrasts using the RMA procedure provided in the *affy* (v1.40.0) R package.

**Obtaining additional microarray data**

We obtained data for *daf-16* mutants (specifically, for *daf-2(e1370)* vs. *daf-16(mu86);daf-2(e1370)*, as DAF-16 is known to be active in *daf-2(-)* mutants) directly from the authors of Shaw, *et al.* (2007)^12^. To obtain microarray data for *nhr-23(RNAi)*, we downloaded gene fold changes for the GEO series GSE32031, which contains three control samples and three *nhr-23(RNAi)* samples; gene fold changes were calculated using the GEO2R web service (<http://www.ncbi.nlm.nih.gov/geo/geo2r/>). Data for *daf-12*, *hlh-30*, and *lin-14* mutants were obtained similarly from GEO series GSE25633, GSE15762, and GSE8520, respectively. The five recent *C. elegans* Affymetrix experiments (platform GPL200) optained from GEO and used to generate Figure 4a were GSE84894, GSE87052, GSE94704, GSE95636, and GSE100814.

**Generation of artificial neural network for independent component partitioning**

To create an artificial neural network for use in partitioning independent components, we first generated simulated data to use as training, validation and test sets. We generated this data by first randomly permuting the expression values of 100 arrays comprising our *C. elegans* microarray compendium column-wise to create a background devoid of non-random signal, but with a similar gene expression value distribution to real data.

Into this random background we inserted simulated gene modules by first picking a gene to use as a module seed pattern, then changing the expression values of other genes such that they positively or negatively correlated with the expression values of this gene across all or a subset of arrays. We varied the number of genes comprising the simulated module, the strength of adherence of each gene to the seed pattern, the fraction of genes within the module with positive and negative correlation to the seed pattern, and the number of arrays in which this correlation existed. In all, we generated over 10,000 random modules and inserted them into separate sets of random background arrays, so that each array set would contain a single non-random module.

We then attempted to recover each simulated module using ICA. We extracted a single component from each simulated array set and deemed the extraction successful if 3 of the top 5 most strongly weighted genes in this component were in the simulated module. For successful extractions, we calculated the optimal partitioning thresholds for the positive and negative hemi-modules, as well as the skewness and kurtosis of the module definition vector using the *moments* (v0.13) R package (<http://CRAN.R-project.org/package=moments>).

Using this data, we trained an artificial network to predict the optimal partitioning thresholds for an independent component from the skewness and kurtosis of its gene weights using the *neuralnet* (v1.32) R package (<http://CRAN.R-project.org/package=neuralnet>). We generated another simulated module set in the same manner as the first to use as a validation set, and varied the architecture of the artificial neural network until the prediction performance reached a maximum value. This occurred when the artificial neural network contained two hidden layers (Supplementary Figure 7), each with 11 nodes (Supplementary Figure 8). We confirmed that the artificial neural network was not over-fit to the validation set by measuring its performance in a third set of simulated data, the test set. Performance on this set was similar to that on the validation set. The structure of this artificial neural network is shown in Supplementary Figure 8).

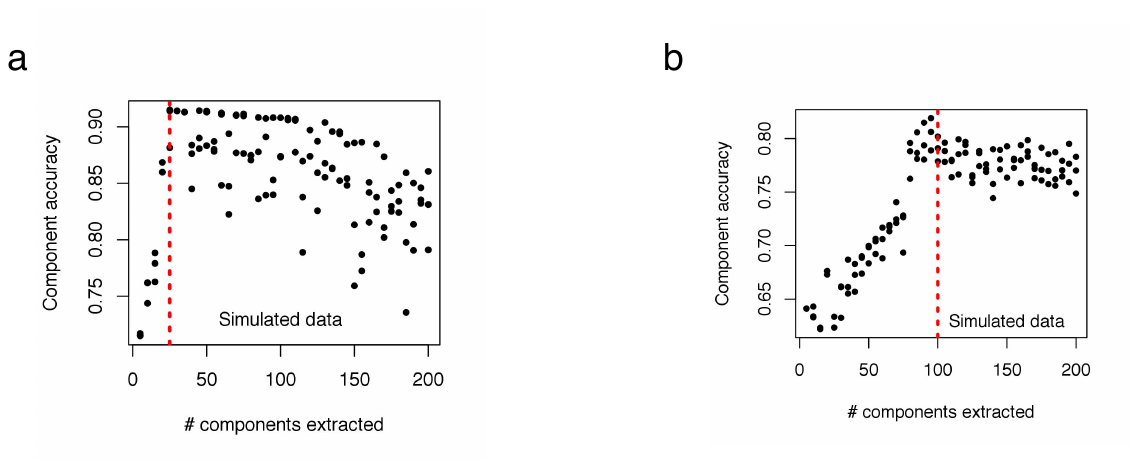

**Supplementary Figure 1. Module extraction using ICA on simulated data.** Results using simulated data show that increasing the number of extracted components improves the accuracy of modules predicted using ICA until this value matches the true number of modules (red dashed line), after which it declines. (**A**) 25 simulated modules, (**B**) 100 simulated modules. Accuracy is defined as average maximum Pearson correlation between extracted and simulated modules.

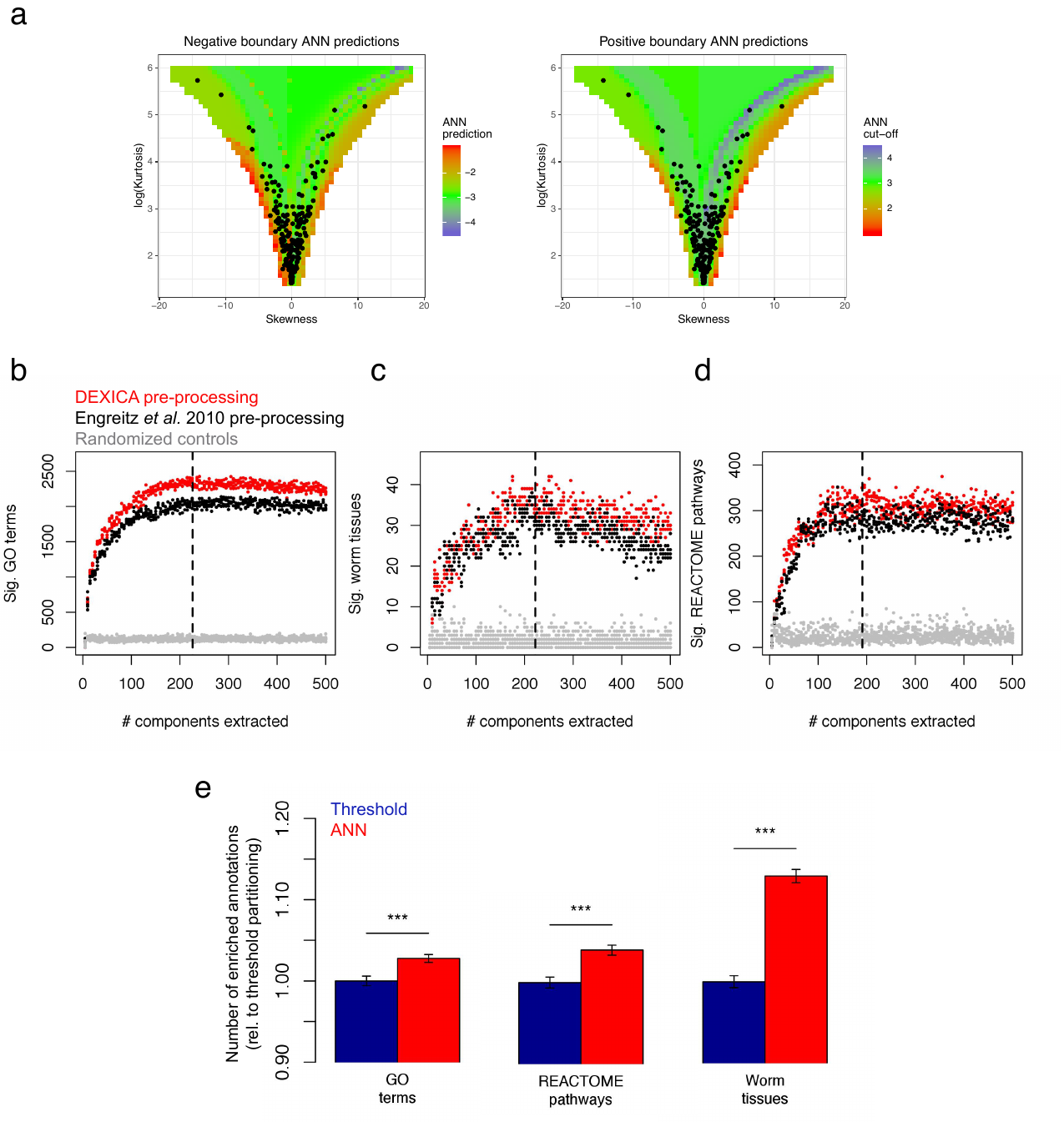

**Supplementary Figure 2. Comparison of a fixed threshold approach and an artificial neural network (ANN) approach for module discretization.** When searching for optimal parameters for gene module prediction (see Figure 2), we applied two different module-partitioning methods. Figure 2 shows results using the ANN-based approach we developed, and (**A**) shows thresholds predicted by the ANN for various skewness and kurtosis combinations, as well as where our predicted hemi-modules (black points) lie in this space . (**B**)-(**D**) show results using the fixed-threshold method (threshold = +/- 3) for generating discrete sets of genes. As in Figure 2A-C, black points show results from a compendium produced using a preprocessing procedure used by Engreitz et *al.* ^13^; red points show results for the best alternative preprocessing method that we tested; dashed lines indicate the point on the x-axis of each graph at which loess regression curves showed the greatest difference between red points and results from randomized control modules (grey points). (**E**) shows the mean performance of ANN discretization relative to fixed threshold discretization for all data points in A-C and Figure 2A-C. Error bars indicate SEM; all comparisons are statistically significant (*** = p < 2.2e-16).

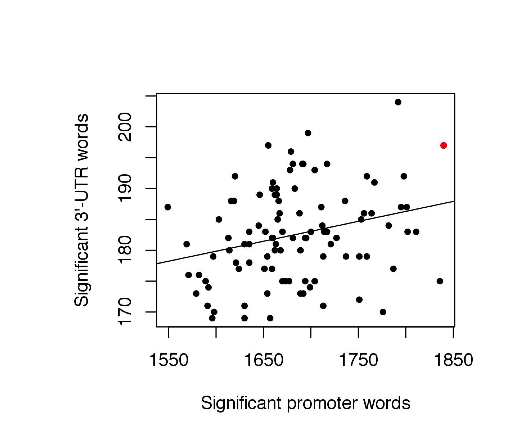

**Supplementary Figure 3. Significant words in predicted gene modules.** We generated 100 sets of gene modules from the *C. elegans* microarray compendium using the optimal parameters indicated by the tests in Figures 2A-C and Supplementary Figures 2B-D. Among these module sets, there is a significant correlation between the total number of promoter words that are significant in at least one module and the number of 3’-UTR words that are significant in at least one module (R = 0.27, p = 6.5e-3). As our final *C. elegans* module set, we chose the set with the best mean rank in these two criteria (indicated by red point in figure).

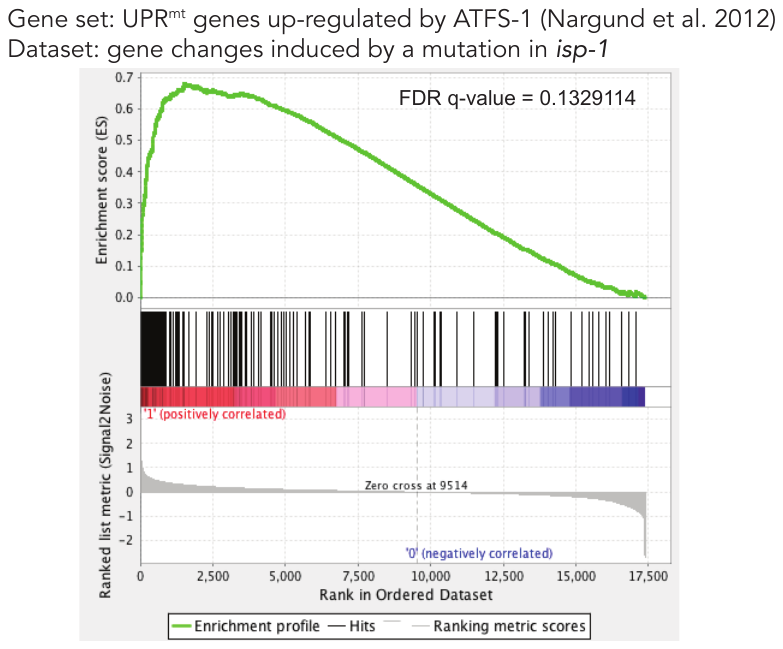

**Supplementary Figure 4. GSEA analysis of *isp-1* gene expression changes and ATFS-1 regulated genes identified in Nargund *et al*. 2012.** Leading edge analysis^3^ shows that genes at the top of the ranked list of genes differentially expressed in *isp-1* mutants are enriched for the ATFS-1 gene set from Nargund *et al*. 2012^14^ (this gene set includes 391 genes). However, the enrichment is not statistically significant.

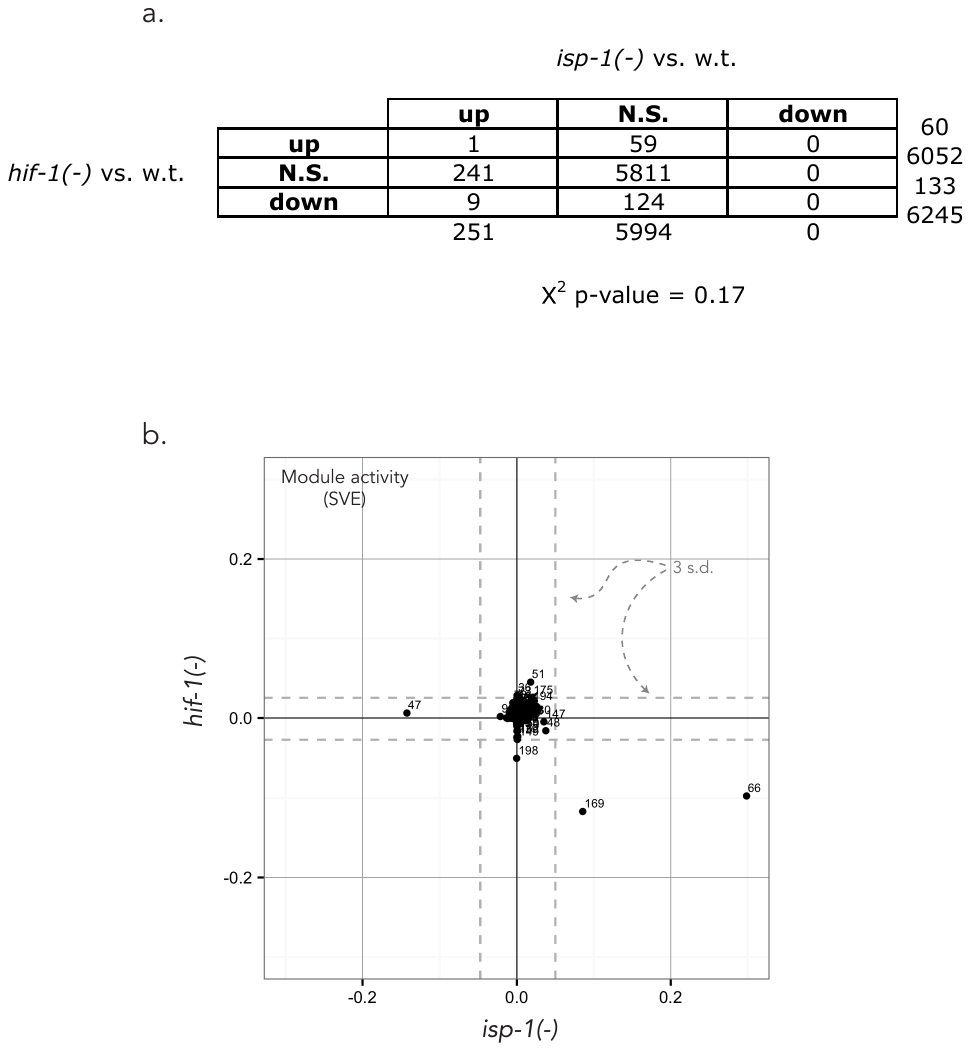

**Supplementary Figure 5. Contingency table for *isp-1(-)* vs. *hif-1(-)* microarray comparison.** (**A**) The *isp-1(-)* and *hif-1(-)* datasets we used in this study were conducted on different microarray platforms, and both datasets contained a substantial number of flagged data points (excluded due to low quality), such that the number of complete pair-wise observations between the two sets was only 6245. The table shows the overlap between the number of significantly up-regulated, significantly down-regulated, and non-significant genes in the two datasets. The *X^2^* p-value for this table is 0.17; thus, there is not a significant degree of overlap among the significant genes of these two datasets. (**B**) Modules 66 and 169 are highly active under both *isp-1*(-) and *hif-1*(-) conditions and are active in opposite directions, consistent with activity on HIF-1 in *isp-1* mutants.

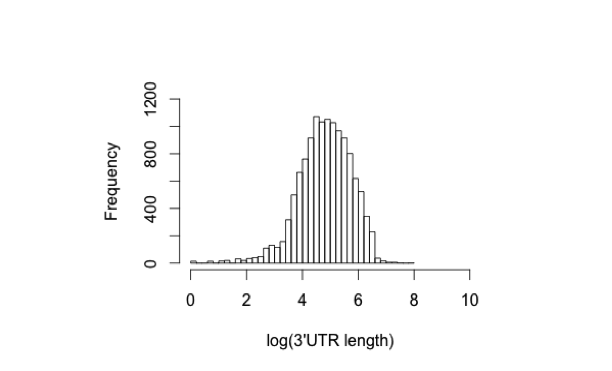

**Supplementary Figure 6. Distribution of *C. elegans* 3’-UTR lengths.** The distribution of *C. elegans* 3’-UTR lengths is approximately log-normal.

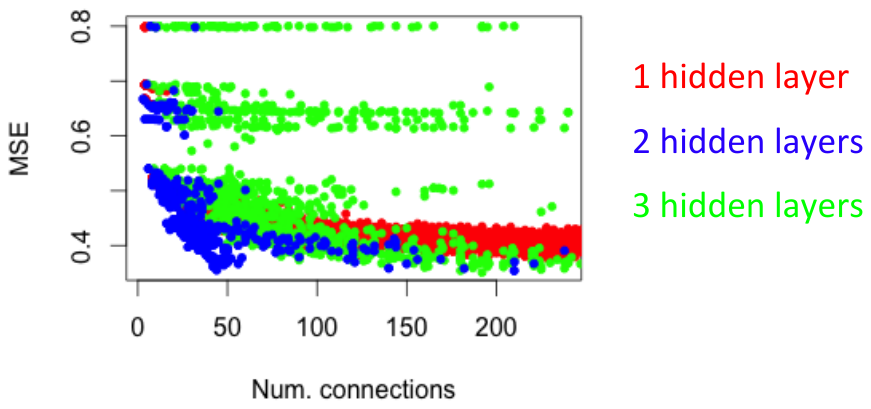

**Supplementary Figure 7. Optimization of artificial neural network for module partitioning.** We tested the effect of various parameters on the prediction performance of artificial neural network discretization of simulated modules. Shown here are the results of varying the number of hidden layers in the network and the number of nodes in each layer; the x-axis shows the total number of connections in the network, the y-axis shows the mean squared error of each network. Colors indicate the number of hidden layers.

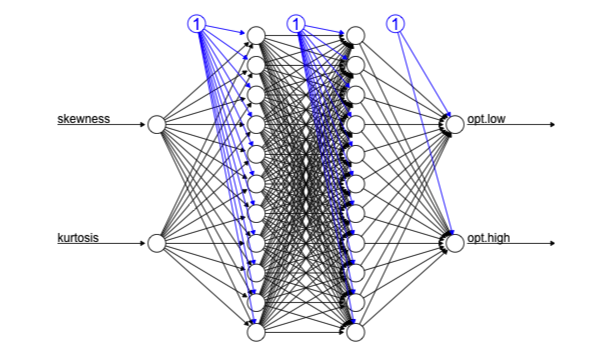

**Supplementary Figure 8. Schematic of optimized artificial neural network for independent component discretization.** The figure shows the structure of the best performing ANN generated by our optimization tests. The skewness and kurtosis of the source matrix weights (columns of the S matrix) of an independent component are used as input to the network, and the predicted optimal discretization thresholds are generated as output.

**Supplementary Tables**

Provided as separate files

**Supplementary Table 1. Genes that belong to each of the 209 DEXICA modules (i.e. partitioned S matrix).** The two hemi-modules are designated by opposite signs.

**Supplementary Table 2. Summary of module annotations**. For each module, the perturbation (from the 716 perturbations in the microarray compendium) in which that module explains the greatest amount of variance (i.e. the perturbation that activates that module the most strongly) is indicated. “None” indicates that a module does not explain more than 5% of the variance in any perturbation (SVE is between -0.05 and 0.05). The top two GO terms from each GO category that are significantly enriched in each hemi-module are also indicated. In cases where enriched GO terms and the perturbation show clear semantic agreement, or if we were able to assign a consensus annotation to a module by other means, this is indicated as a “process most likely represented by the module”.

**Supplementary Table 3. Comparison of module activity in all possible pairs of perturbations (contrasts) in the microarray compendium.** Pairs are ranked by the absolute value of the Pearson correlation coefficient calculated based on activity of each module (SVE).

**Supplementary Table 4**. **Module-weighted GO annotations of each gene**. Weighted associations between each gene and each GO term are shown. Numbers represent z-scores of gene weights in each GO category. Only terms associated with at least 15.

**Supplementary references**

1.	The ENCODE (ENCyclopedia Of DNA Elements) Project. *Science* **306,** 636–640 (2004).

2.	Gerstein, M. B. *et al.* Integrative analysis of the Caenorhabditis elegans genome by the modENCODE project. *Science* **330,** 1775–1787 (2010).

3.	Subramanian, A. *et al.* Gene set enrichment analysis: a knowledge-based approach for interpreting genome-wide expression profiles. *PNAS* **102,** 15545–15550 (2005).

4.	Kim, S. Y. & Volsky, D. J. PAGE: parametric analysis of gene set enrichment. *BMC Bioinformatics* **6,** 144 (2005).

5.	Luo, W., Friedman, M. S., Shedden, K., Hankenson, K. D. & Woolf, P. J. GAGE: generally applicable gene set enrichment for pathway analysis. *BMC Bioinformatics* **10,** 161 (2009).

6.	Gatti, D. M., Barry, W. T., Nobel, A. B., Rusyn, I. & Wright, F. A. Heading down the wrong pathway: on the influence of correlation within gene sets. *BMC Genomics* **11,** 574 (2010).

7.	Goeman, J. J. & Buhlmann, P. Analyzing gene expression data in terms of gene sets: methodological issues. *Bioinformatics* **23,** 980–987 (2007).

8.	Tamayo, P., Steinhardt, G., Liberzon, A. & Mesirov, J. P. The limitations of simple gene set enrichment analysis assuming gene independence. *Stat Methods Med Res* **25,** 472–487 (2016).

9.	Bussemaker, H. J., Li, H. & Siggia, E. D. Building a dictionary for genomes: identification of presumptive regulatory sites by statistical analysis. *PNAS* **97,** 10096–10100 (2000).

10.	Durinck, S., Spellman, P. T., Birney, E. & Huber, W. Mapping identifiers for the integration of genomic datasets with the R/Bioconductor package biomaRt. *Nat Protoc* **4,** 1184–1191 (2009).

11.	van Helden, J., Andre, B. & Collado-Vides, J. Extracting regulatory sites from the upstream region of yeast genes by computational analysis of oligonucleotide frequencies. *J Mol Biol* **281,** 827–842 (1998).

12.	Shaw, W. M., Luo, S., Landis, J., Ashraf, J. & Murphy, C. T. The C. elegans TGF-beta Dauer pathway regulates longevity via insulin signaling. *Current biology : CB* **17,** 1635–1645 (2007).

13.	Engreitz, J. M., Daigle, B. J., Marshall, J. J. & Altman, R. B. Independent component analysis: mining microarray data for fundamental human gene expression modules. *J Biomed Inform* **43,** 932–944 (2010).

14.	Nargund, A. M., Pellegrino, M. W., Fiorese, C. J., Baker, B. M. & Haynes, C. M. Mitochondrial import efficiency of ATFS-1 regulates mitochondrial UPR activation. *Science* **337,** 587–590 (2012).
